# Flexible coordinator and switcher hubs for adaptive task control

**DOI:** 10.1101/822213

**Authors:** Carrisa V Cocuzza, Takuya Ito, Douglas Schultz, Danielle S Bassett, Michael W Cole

**Affiliations:** Center for Molecular and Behavioral Neuroscience, Rutgers University, Newark, NJ, USA, 07102; Behavioral and Neural Sciences PhD Program, Rutgers University, Newark, NJ, USA, 07102; Center for Brain Biology and Behavior, University of Nebraska-Lincoln, Lincoln, NE, USA, 68588; Departments of Bioengineering, Physics & Astronomy, Electrical & Systems Engineering, and Neurology, University of Pennsylvania, Philadelphia, PA, USA, 19104

**Keywords:** cognitive control, network dynamics, executive function, network interactions, cognitive flexibility, task representation

## Abstract

Functional connectivity studies have identified at least two large-scale neural systems that constitute cognitive control networks – the frontoparietal network (FPN) and cingulo-opercular network (CON). Control networks are thought to support goal-directed cognition and behavior. It was previously shown that the FPN flexibly shifts its global connectivity pattern according to task goal, consistent with a “flexible hub” mechanism for cognitive control. Our aim was to build on this finding to develop a functional cartography (a multi-metric profile) of control networks in terms of dynamic network properties. We quantified network properties in (male and female) humans using a high-control-demand cognitive paradigm involving switching among 64 task sets. We hypothesized that cognitive control is enacted by the FPN and CON via distinct but complementary roles reflected in network dynamics. Consistent with a flexible “coordinator” mechanism, FPN connections were varied across tasks, while maintaining within-network connectivity to aid cross-region coordination. Consistent with a flexible “switcher” mechanism, CON regions switched to other networks in a task-dependent manner, driven primarily by reduced within-network connections to other CON regions. This pattern of results suggests FPN acts as a dynamic, global coordinator of goal-relevant information, while CON transiently disbands to lend processing resources to other goal-relevant networks. This cartography of network dynamics reveals a dissociation between two prominent cognitive control networks, suggesting complementary mechanisms underlying goal-directed cognition.

**Significance Statement:** Cognitive control supports a variety of behaviors requiring flexible cognition, such as rapidly switching between tasks. Furthermore, cognitive control is negatively impacted in a variety of mental illnesses. We used tools from network science to characterize the implementation of cognitive control by large-scale brain systems. This revealed that two systems – the frontoparietal (FPN) and cingulo-opercular (CON) networks – have distinct but complementary roles in controlling global network reconfigurations. The FPN exhibited properties of a flexible coordinator (orchestrating task changes), while CON acted as a flexible switcher (switching specific regions to other systems to lend processing resources). These findings reveal an underlying distinction in cognitive processes that may be applicable to clinical, educational, and machine learning work targeting cognitive flexibility.

## Introduction

Theories of cognitive control – processes supporting goal-directed cognition and behavior – suggest the need for flexibly reconfigurable neural systems to support controlled processing (Desimone and Duncan, 1995; Miller and Cohen, 2001; Schneider and Chein, 2003; Cole et al., 2013b). In order for an individual’s goals to be implemented, goal-relevant information must be appropriately represented across large-scale neural systems, or networks. Importantly, goals and goal-relevant information are subject to change over time (such as sensorimotor information that corresponds to changing task conditions). Processing these dynamic changes must be guided amongst neural systems that represent goal-relevant information. Cognitive control networks are proposed to enact this guidance via network interactions that are flexible with respect to the current task context (Waskom et al., 2014). Thus, we focus here on the role of large-scale network dynamics as task goals are updated across 64 systematically-related task contexts (Fig. 1).

**Figure 1.**
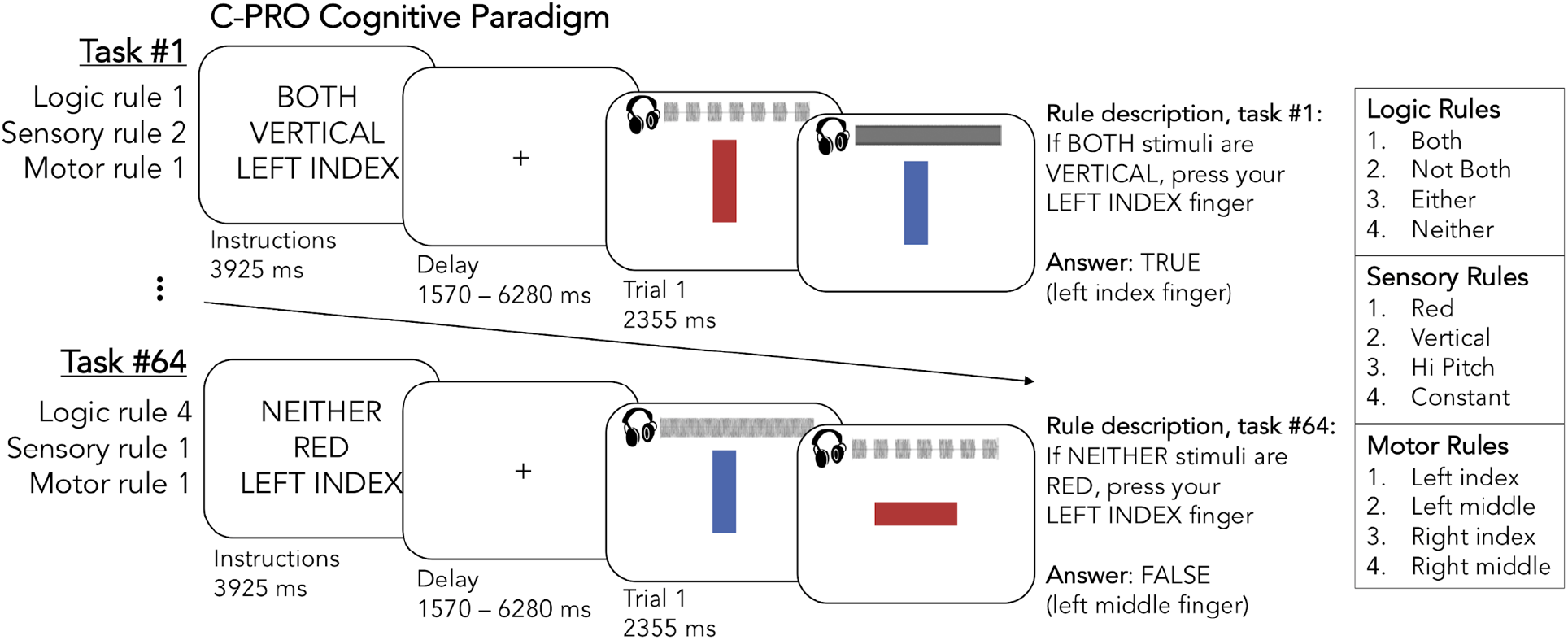
The Concrete Permuted Rule Operations (C-PRO) cognitive paradigm. First, an instruction screen presented the rules for a given task (3925 ms). Participants next applied these rules to pairs of consecutively presented audiovisual stimuli (auditory waveforms are visually depicted here, but were only presented audibly to participants). Two example task-rule sets are depicted, as well as how participants were trained to interpret the rules (e.g., rule descriptions on the right-most portion of the figure) (see Materials and Methods for details). The 12 possible rules are listed on the right.

The theoretical insight that large-scale network interactions are essential to cognitive control evolved over the last several decades, beginning with empirical observations (e.g., Fuster et al., 1985), which then led to the biased competition theory (Desimone et al., 1990, 1995). This theory focused on lateral prefrontal cortex influencing the visual system by biasing its competition for attentional resources toward goal-relevant representations. Building on the biased competition theory, the guided activation theory generalized this prefrontal network mechanism to all task domains. This theory proposed a general role for top-down prefrontal influences in accomplishing task goals (Miller and Cohen, 2001). More recently, the flexible hub theory generalized the guided activation theory beyond prefrontal cortex to the entire frontoparietal network (FPN) and formalized the importance of cross-network, global connectivity changes in implementing cognitive control (Cole et al., 2013b). The present study builds on this work to further verify and expand the flexible hub theory.

Simultaneous with these advances in theory have been observations of a second major neural system supporting cognitive control: the cingulo-opercular network (CON). Like the FPN, the CON is active as a function of cognitive control demands across a wide variety of tasks (Dosenbach et al., 2006; Yeo et al., 2015; Crittenden et al., 2016). However, CON and FPN are not equally active for all task conditions (Dosenbach et al., 2006; Yeo et al., 2015) and they maintain distinct functional network architectures in terms of resting-state functional connectivity (rsFC) (Dosenbach et al., 2007; Power et al., 2011; Ji et al., 2019) and task-state functional connectivity (tFC) (Cole et al., 2014; Crittenden et al., 2016). Moreover, the specific functional contributions of CON regions have not been fully established, with some studies suggesting that CON regions specify overall task set modes of processing (Dosenbach et al., 2007; Sadaghiani and D’Esposito, 2015), and others emphasizing the CON’s role in reactive (phasic) attention (Seeley et al., 2007), and relatedly, conflict processing (Cole et al., 2009; Botvinick, 2007; Braem et al., 2019). Ultimately, unlike the FPN, the relationship between the CON and the flexible hub theory (and the theories it builds upon) remains unclear.

The present study builds on our prior work demonstrating flexible hub properties in FPN regions (Cole et al., 2013b), expanding on the characterization of these FPN network mechanisms, while also investigating CON network mechanisms. We previously found that FPN’s global tFC patterns flexibly updated according to task demands more than any other network, including CON (Cole et al., 2013b). However, given that large-scale network dynamics are central to cognitive control, and given that both the CON and FPN contain hubs (Power et al., 2011; Ito et al., 2017), we hypothesized that CON reflects flexible hub properties in addition to FPN. Unlike FPN’s continuous goal-coordinating role, we expected CON to exhibit a more discrete network switching mechanism, reflecting its proposed role in specifying overall task-set modes of processing (Dosenbach et al., 2007; Sadaghiani and D’Esposito, 2015). Consistent with this, we found that FPN regions act as “flexible coordinators” and CON regions as “flexible switchers”, providing separate but complementary network mechanisms in support of cognitive control (Fig. 2).

**Figure 2.**
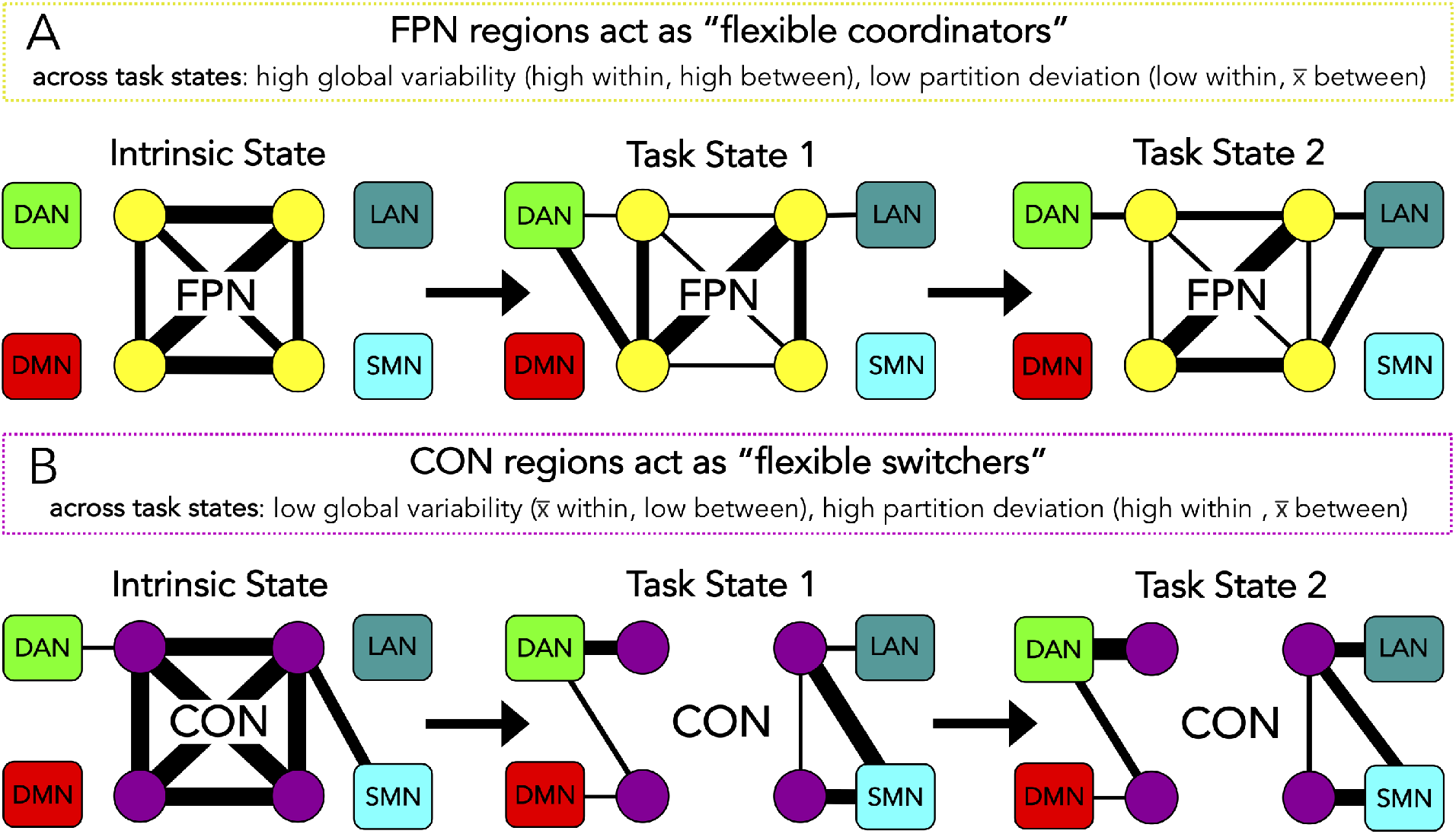
Schematic depictions of cognitive control network functional properties. (See Materials and Methods and Results for details on network measures). In each panel, a “toy” version of the control network is prominently depicted in the center (with a reduced number of regions, or nodes, and simplified within-network connections), and out-of-network exemplars are depicted as truncated and surrounding the control network of interest *(DAN:* dorsal attention network, *LAN:* language network, *DMN:* default mode network, and *SMN:* somatomotor network). Each of these surrounding networks also contains within-network regions and connections, but these were not depicted here for simplicity. *(**A**)* Regions in the frontoparietal network (FPN) acted as flexible coordinators. This entailed high global variability (GVC) and low partition deviation across task states. From example task state one to task state two, FPN regions maintained their within-network connectivity (low deviation) and out-of-network connectivity changes were variable across states (high GVC). (See Results for details). (***B***) Regions in the cingulo-opercular network (CON) acted as flexible switchers. This entailed low global variability and high partition deviation. From example task state one to task state two, CON regions dropped their within-network connectivity (high deviation) and out-of-network connectivity changes were consistent across states (low GVC). (See Results for details).

## Materials and Methods

### Participants

Right-handed, healthy adult participants (*N* = 106) were recruited from Rutgers University and the surrounding Newark, New Jersey community. Six participants were excluded from analyses due to technical errors, leaving a total sample size of *N* = 100 (Table 1 and Table 2 detail demographic characteristics). To improve replicability, we used a split-sample validation approach (Anderson and Magruder, 2017) with a random subset of *n* = 50 comprising a discovery dataset (Table 1), and the remaining *n* = 50 comprising a replication dataset (Table 2). All participants provided informed consent in accordance with protocols approved by the Institutional Review Board of Rutgers University-Newark. Each participant provided or completed the following: (1) demographic information and intake survey questions, (2) the National Institutes of Health Cognition Toolbox (Gershon et al., 2013), including a neuropsychological battery, (3) behavioral training on the C-PRO task (outside the scanner), (4) resting-state fMRI, and (5) C-PRO task fMRI. The subsets of data assessed herein included (1), (4), and (5) (see Schultz et al., 2019 for assessment of other variables). As listed in the right-most columns of Tables 1 and 2, there were no significant differences between identified genders on the distributions of age, ethnicity, or education.

**Table 1.** Demographic characteristics of the discovery dataset *(n* = 50). There were no significant differences between identified genders on the distributions of age, ethnicity, or education (right column). *The measure of center used for the age variable was the mean, and for categorical variables of ethnicity/race and education it was the mode. For the education variable, student refers to “some college”. **Hypothesis testing of significant differences between males and females. Age: two-sample t-test adjusted for unequal sample sizes. Ethnicity/race and education: a chi-square test of independence.

| Table 1. Demographic characteristics of the discovery dataset ($n=50$). | | | | | | | | | |
| --- | --- | --- | --- | --- | --- | --- | --- | --- | --- |
| | Male ($n=19$) | | | | Female ($n=31$) | | | | |
|  | n | % | center* | SD (+/-) | n | % | center* | SD (+/-) | test (male vs female)** |
| Age (years) | | | 21.2 | 2.9 | | | 19.9 | 1.6 | $t(48)=1.71, p=0.10$ |
| 18-24 | 17 | 89.5 |  |  | 31 | 100 |  |  |  |
| 25-34 | 2 | 10.5 |  |  | 0 | 0 |  |  |  |
| 35-44 | 0 | 0 |  |  | 0 | 0 |  |  |  |
| Ethnicity/Race | | | White | n/a | | | Black | n/a | $\chi^2(5, N=50)=6.29, p=0.18$ |
| American Indian or Alaskan Native | 0 | 0 |  |  | 0 | 0 |  |  |  |
| Asian | 5 | 26.3 |  |  | 10 | 32.3 |  |  |  |
| Black or African American | 3 | 15.8 |  |  | 12 | 38.7 |  |  |  |
| Hispanic or Latino | 4 | 21.1 |  |  | 2 | 6.5 |  |  |  |
| Native Hawaiian or Pacific Islander | 0 | 0 |  |  | 0 | 0 |  |  |  |
| White or Caucasian | 7 | 36.8 |  |  | 6 | 19.4 |  |  |  |
| Other | 0 | 0 |  |  | 1 | 3.2 |  |  |  |
| Education | | | Student | n/a | | | Student | n/a | $\chi^2(2, N=50)=1.27, p=0.26$ |
| Some college/Associate's degree | 15 | 79 |  |  | 28 | 90.3 |  |  |  |
| College/Graduate degree | 4 | 21.1 |  |  | 3 | 9.7 |  |  |  |
| Not reported | 0 | 0 |  |  | 0 | 0 |  |  |  |

**Table 2.** Demographic characteristics of the replication dataset (*n* = 50. All table features are the same as in Table 1. Note that there were no significant differences between identified genders on the distributions of age, ethnicity, or education (right column).

| Table 2. Demographic characteristics of the replication dataset (n=50). |  |  |  |  |  |  |  |  |  |
| --- | --- | --- | --- | --- | --- | --- | --- | --- | --- |
|  | Male (n=25) |  |  |  | Female (n=25) |  |  |  |  |
|  | n | % | center* | SD (+/-) | n | % | center* | SD (+/-) | test (male vs female)** |
| Age (years) |  |  | 25 | 5.4 |  |  | 23.2 | 3.5 | t(48)=1.44, p=0.16 |
| 18-24 | 12 | 48 |  |  | 17 | 68 |  |  |  |
| 25-34 | 10 | 40 |  |  | 8 | 32 |  |  |  |
| 35-44 | 3 | 12 |  |  | 0 | 0 |  |  |  |
| Ethnicity/Race | | | Asian | n/a | | | Asian | n/a | $\chi^2(5, N=50)=7.23, p=0.3$ |
| American Indian or Alaskan Native | 3 | 12 |  |  | 0 | 0 |  |  |  |
| Asian | 8 | 32 |  |  | 10 | 40 |  |  |  |
| Black or African American | 5 | 20 |  |  | 3 | 12 |  |  |  |
| Hispanic or Latino | 1 | 4 |  |  | 4 | 16 |  |  |  |
| Native Hawaiian or Pacific Islander | 0 | 0 |  |  | 0 | 0 |  |  |  |
| White or Caucasian | 7 | 28 |  |  | 8 | 32 |  |  |  |
| Other | 1 | 4 |  |  | 0 | 0 |  |  |  |
| Education | | | Graduate | n/a | | | Student | n/a | $\chi^2(2, N=50)=1.47, p=0.48$ |
| Some college/Associate's degree | 9 | 36 |  |  | 12 | 48 |  |  |  |
| College/Graduate degree | 13 | 52 |  |  | 12 | 48 |  |  |  |
| Not reported | 3 | 12 |  |  | 1 | 4 |  |  |  |

### Concrete Permuted Rule Operations (C-PRO) paradigm

The C-PRO paradigm was designed to involve rapid instructed task learning (RITL) through compositionally combining various task rules (Cole et al., 2010, 2013a; Ito et al., 2017). This further provided a high demand on cognitive control across all C-PRO task states. We used a modified version of the PRO paradigm from Cole et al. (2010), which was previously introduced by Ito et al. (2017) (Fig. 1). This paradigm permutes rules across three domains: four logic rules (both, not both, either, and neither), four sensory rules (red color, vertical orientation, high pitch sound, and constant tone), and four motor rules (left index, right index, left middle, and right middle fingers). This amounts to 12 rule sets represented 16 times across 64 unique task states. The software for presenting the task was E-Prime version 2.0.10.353 (Schneider et al., 2002).

In each task state, an initial instruction screen was presented for 3925 ms for participants to memorize a given permuted rule set (Fig. 1). This was followed by a jittered delay (1570 – 6280 ms, randomized from a uniform distribution), then three trials of paired audiovisual stimuli for participants to adjudicate based upon the given rule set (2355 ms each trial; inter-trial interval of 1570 ms). Another jittered delay occurred at the end of each task state (7850 – 12560 ms, randomized), which was immediately followed by the next permuted rule set instruction screen. An example instruction screen (Fig. 1; task state one) read: “BOTH, VERTICAL, LEFT INDEX”, indicating: “If both stimuli are vertical, press your left index finger”. In each of the three trials that followed, participants judged if both paired stimuli were vertically oriented, and either pressed the left index finger button to indicate “true” or the left middle finger button to indicate “false” (a judgement of “false” was always the same hand but opposite finger). Importantly, stimuli were always presented with auditory and visual features concurrently. Thus, focusing on the sensory rule given by the instructions was paramount (i.e., “VERTICAL” indicated that one should ignore auditory information, color information, and only focus on line orientation). Additionally, participants were required to remember and apply conditional logic and nontrivial motor commands each trial. Altogether this multitask behavioral paradigm is condition-rich and necessitates ongoing cognitive control.

Each participant completed a training session outside the scanner and a testing session within the scanner (task-state fMRI) 30 minutes later. During the training session participants equally practiced four rule sets that contained all 12 rules. This practice set was counterbalanced amongst participants and supplementary instruction was provided for training purposes (e.g., use the same hand but opposite finger to indicate “false”). Task fMRI scans were performed in eight runs, altogether containing 64 task state miniblocks, twice over (e.g., 128 task miniblocks), with each block composed of a permuted rule set (Fig. 1). Each task fMRI run was approximately eight minutes in duration, and identical miniblocks were never presented consecutively. Overall, mean performance was 83.47% correct (*SD* = 9.00%). There was no significant difference in performance (percent correct) between males (*M* = 83.65%, *SD* = 10.44%) and females (*M* = 83.33%, *SD* = 7.80%); *t*(74.71) = 0.17, *p* = 0.87.

### Experimental design and statistical analysis

Participants were randomly allocated to either a discovery dataset (*n* = 50) or replication dataset (*n* = 50) (Table 1 and Table 2, respectively). The replication dataset was not analyzed until after analyses of the discovery dataset were complete. Analyses of replication data were identical to analyses of discovery data (using the same code, including all chosen parameters), and additionally included measures of similarity between replication and discovery results to quantify expected generalizability (Anderson and Magruder, 2017).

Whenever multiple comparisons were addressed, we utilized the Max-T nonparametric permutation testing approach (10,000 permutations unless otherwise specified) with maxima-derived 95% confidence intervals for statistical hypothesis testing against zero (Blair and Karniski, 1993; Nichols and Holmes, 2002). To analyze the similarity of two correlation (weighted adjacency) matrices we used the Mantel permutation test, which performs a Pearson’s correlation across the upper triangles (off-diagonal) of the matrices (Mantel, 1967; Glerean et al., 2016). The Mantel test is more conservative than a standard comparison between connectivity matrices because it takes into account the fact that observations in distance/similarity matrices are not independent (an assumption of both parametric and standard non-parametric tests). In each Mantel analysis, we again used nonparametric permutation procedures to derive statistics that make minimal assumptions about probability distribution (10,000 permutations unless otherwise specified). Henceforth we will describe these matrix similarity statistics as Mantel-r.

### MRI parameters

All MRI data were collected at the Rutgers University Brain Imaging Center (RUBIC). When possible, the best practices suggested by the Human Connectome Project preprocessing pipelines were followed (Glasser et al., 2013). A 3T, 32-channel head coil within a Siemens Trio scanner was used to obtain multiband, whole-brain, and echo-planar imaging (EPI). The repetition time (TR) was 785 ms; the echo time (TE) was 34.8 ms; the flip angle was 55°; the bandwidth was 1924 Hz/Px; the in-plane field-of-view (FoV) read was 208 mm; 72 slices; 2.0 mm isotropic voxels; and the multiband acceleration factor was 8. Whole-brain and high-resolution T1-weighted and T2-weighted anatomical scans were also acquired, with an isotropic voxel resolution of 0.8 mm. Spin echo field maps were obtained in both the anterior-posterior and posterior-anterior directions. Resting-state fMRI scans were 14 minutes in duration, amounting to 1070 TRs. Each task (i.e., C-PRO) fMRI run was approximately eight minutes in duration, adding up to approximately one hour in the scanner for the task session (36 TRs per task miniblock; 4608 TRs altogether).

### fMRI preprocessing

The open-source Human Connectome Project minimal preprocessing pipeline (Glasser et al., 2013), version 3.5.0, was applied to all neuroimaging data. This included: anatomical reconstruction and segmentation; EPI reconstruction, segmentation, and spatial normalization to a standard template; intensity normalization; and motion correction. The resulting data was in CIFTI 64k-vertex grayordinate space, and all subsequent analyses were performed in MATLAB R2014b (The Mathworks, Inc.). Following minimal preprocessing, vertices were parcellated into 360 cortical regions (180 per hemisphere) per the Glasser et al. (2016) atlas. To parcellate each of these regions, we calculated the average time series of enclosed vertices.

Next, we performed nuisance regression on parcellated resting-state and task-state data using 6 motion parameters plus their derivatives (totaling 12 motion parameters), and volumetrically-extracted ventricle and white matter time series (via FreeSurfer http://surfer.nmr.mgh.harvard.edu/), plus their first derivatives (16 regressors overall). Note that global signal was not removed due to evidence that it can artificially introduce negative relationships (Murphy et al., 2009). Task time series were further processed to account for confounding effects introduced by simultaneous sensory inputs (e.g., left and right primary visual area, V1) and their downstream effects by fitting a general linear model (GLM) to task activity estimated by a finite impulse response (FIR) function. This removal of cross-event mean task-locked activity has been shown to reduce task-evoked correlation false positives while retaining most (~90%) of the correlated variance between fMRI time series and without inflating false negatives (Cole et al., 2019). In the task GLM, each task run was separately demeaned, and drift was accounted for with a per-run linear trend.

### Functional connectivity estimation

Functional connectivity (FC) was estimated for parcellated (region-wise), pre-processed data, per participant and per state (one resting state and 64 C-PRO task states). Across the whole cortex, we utilized Fisher’s Z-transformed Pearson correlation coefficients to compute interregional relationships of blood-oxygen-level dependent (BOLD) time series, resulting in 360 by 360 connectivity matrices. Given the complex nature of subsequent analyses (i.e., network metrics) we chose this method of FC estimation for simplicity and wide-reaching comprehension. In the present study, connectivity estimates tended to decrease from rest to task, a finding that has been observed across numerous prior studies (that utilized various model species and neural recording methods) (Cohen and Maunsell, 2009; He, 2013; Cole et al., 2014; Ponce-Alvarez et al., 2015) and has well-founded neural mechanisms (Ito et al., 2019).

We chose to use FIR regression to remove cross-block mean task-evoked activations prior to Pearson correlation estimation (sometimes termed “background connectivity”, as in Norman-Haignere et al., 2012) based on recent results demonstrating that this approach was better able to remove confounding effects of task-evoked activity than alternative approaches, such as psychophysiological interactions (PPI) (Cole et al., 2019). Our prior global variability coefficient (see Materials and Methods section below section on network metrics) results were based on generalized PPI connectivity estimates (Cole et al., 2013b), such that the present results provided improved testing of the flexible hub theory.

### Network partition

We applied the cortical portion of the Cole-Anticevic brain-wide network partition (CAB-NP) (Ji et al., 2019; Fig. 3), which was based on publicly available Human Connectome Project data. The CAB-NP was based on resting-state fMRI data across the whole brain, and used the Louvain community detection algorithm to assign parcellated cortical regions (Glasser et al., 2016) into 12 functional networks. The CAB-NP corroborated features of well-known cortical partitions (Gordon et al., 2016; Power et al., 2011; Yeo et al., 2011), yet found novel but robust networks. The CAB-NP was implemented for all analyses except network flexibility, which requires the application of community detection (Louvain Q-modularity; see Materials and Methods section below on network flexibility).

**Figure 3.**
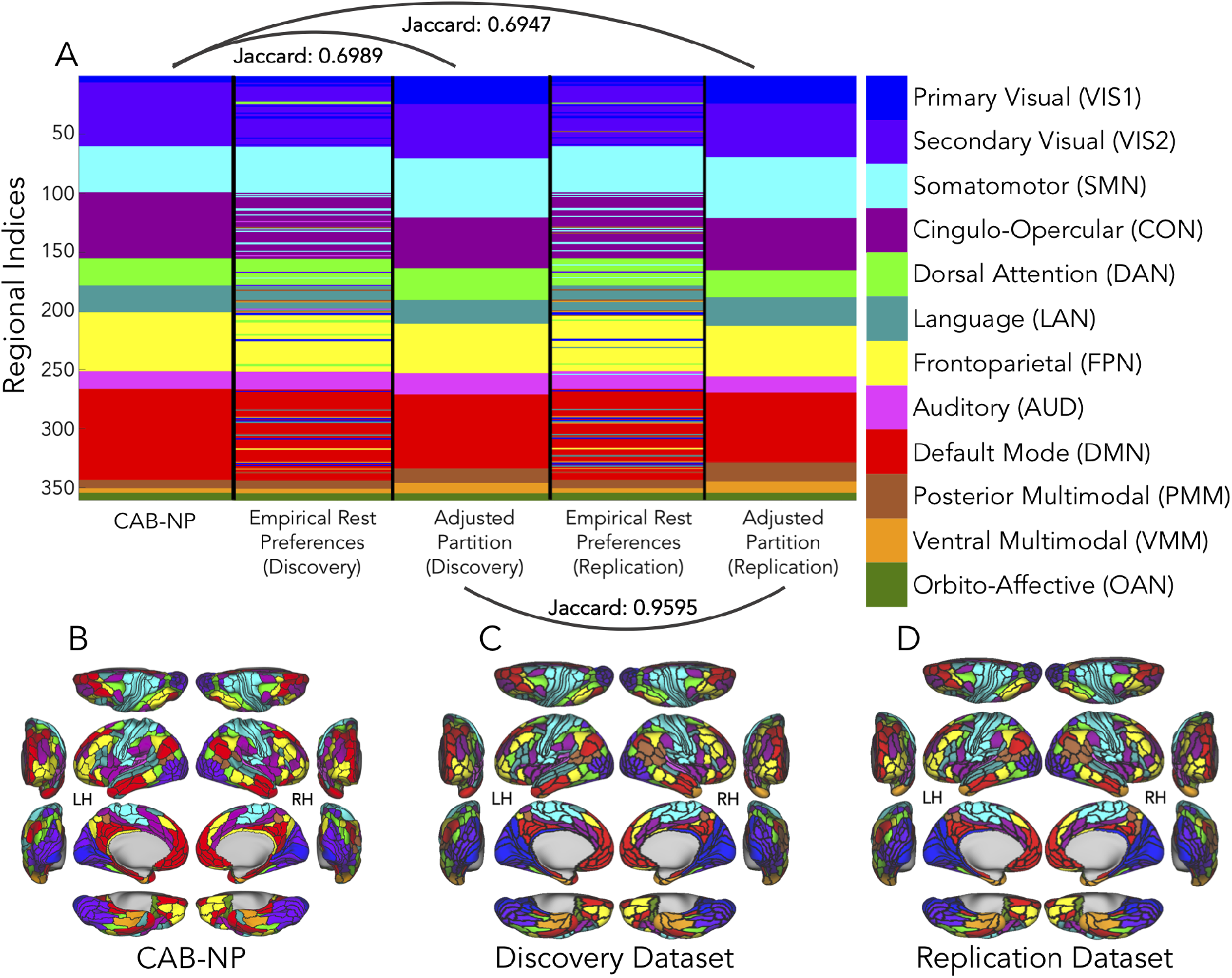
The Cole-Anticevic brain-wide network partition (CAB-NP) adjusted by empirical resting-state FC, for both the discovery and replication datasets. (***A***) Regional (y-axis; Glasser et al., 2016 parcels) assignments are color-coded according to the CAB-NP (rightmost scale). The CAB-NP column depicts the original restingstate network partition by Ji et al., 2019. The empirically-derived rest preferences are shown, unordered, for both the discovery and replication datasets, as well as their ordered counterparts (i.e., “adjusted partitions”). These adjusted partitions were used for all analyses. (***B***) CAB-NP by Ji et al. (2019) projected onto brain regions. (***C***) The empirically-adjusted CAB-NP for the discovery dataset projected onto brain regions. The Jaccard similarity coefficient between the CAB-NP and the empirically-adjusted discovery set partition was 0.6989. (***D***) The empirically-adjusted CAB-NP for the replication dataset projected onto brain regions. The Jaccard similarity coefficient between discovery and replication partitions was 0.9595, suggesting the partition method used herein will have high external validity. The Jaccard similarity coefficient between the CAB-NP and the empirically-adjusted replication set partition was 0.6947. This suggests a relatively high similarity between each of the empirically-adjusted partitions (discovery and replication) and the CAB-NP. The least similarity was observed in the primary visual network, which was expanded to include CAB-NP secondary visual and dorsal attention regions in the empirical adjustments.

Given that our novel network metric (see Materials and Methods section on network partition deviation) quantifies network affiliation changes from an intrinsic partition, it was important (in order to avoid inflated deviation estimates) to ensure that the intrinsic partition was applicable to the present group of subjects. We first partitioned resting-state data by sorting regional FC estimates per the 12 CAB-NP network indices. We then found the maximum FC estimate (i.e., the intrinsic “preference”) for each region (per participant), and tested if its location was equivalent to the CAB-NP. If this index was different from the CAB-NP in over 50% of participants, we reassigned that region to its empirically-derived preference. We henceforth used this empirically-adjusted CAB-NP to sort task-state data into networks (Fig. 3C and 3D).

In select analyses, we probed the similarity of two partitions. To accomplish this, we used the Jaccard index, which is a standard measure of similarity from set theory. For example, the Jaccard index was used to assess the similarity of the empirically-adjusted resting-state partitions of the discovery and replication datasets. We used the MATLAB *jaccard* function, which utilized the “intersection over union” formula on label vectors A and B, with the following equation:

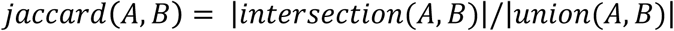

Per state, the intersection equaled the number of true positives (i.e., overlap of two partitions), and the union was the number of true positives summed with the number of false positives and false negatives.

### Network metrics

Interregional connectivity was probed by three network metrics for state-based reconfiguration properties (Medaglia et al., 2015): (1) Global variability coefficient (GVC; Cole et al., 2013b, and relatedly, between-network variability coefficient (BVC; novel but related to Ito et al., 2017), (2) Network flexibility (NF; Bassett et al., 2011, 2013a), and (3) Network partition deviation (deviation; novel). Network metrics were computed across states and averaged across regions that compose a given network, per participant. In analyses that used standardized metrics (i.e., z-scores), standardizations were performed before network averages and standard errors were computed. Figure 4 illustrates the algorithms of these metrics schematically. Table 3 summarizes the primary characteristics of these metrics, including formulae, interpretations, parameterspace considerations, and reliance on a predefined network partition. A predefined network partition is sometimes called a “hard partition”, and refers to the use of a predefined network or community assignment structure, such that each parcellated region is indexed into the partition *a priori* (Sporns and Betzel, 2016).

**Figure 4.**
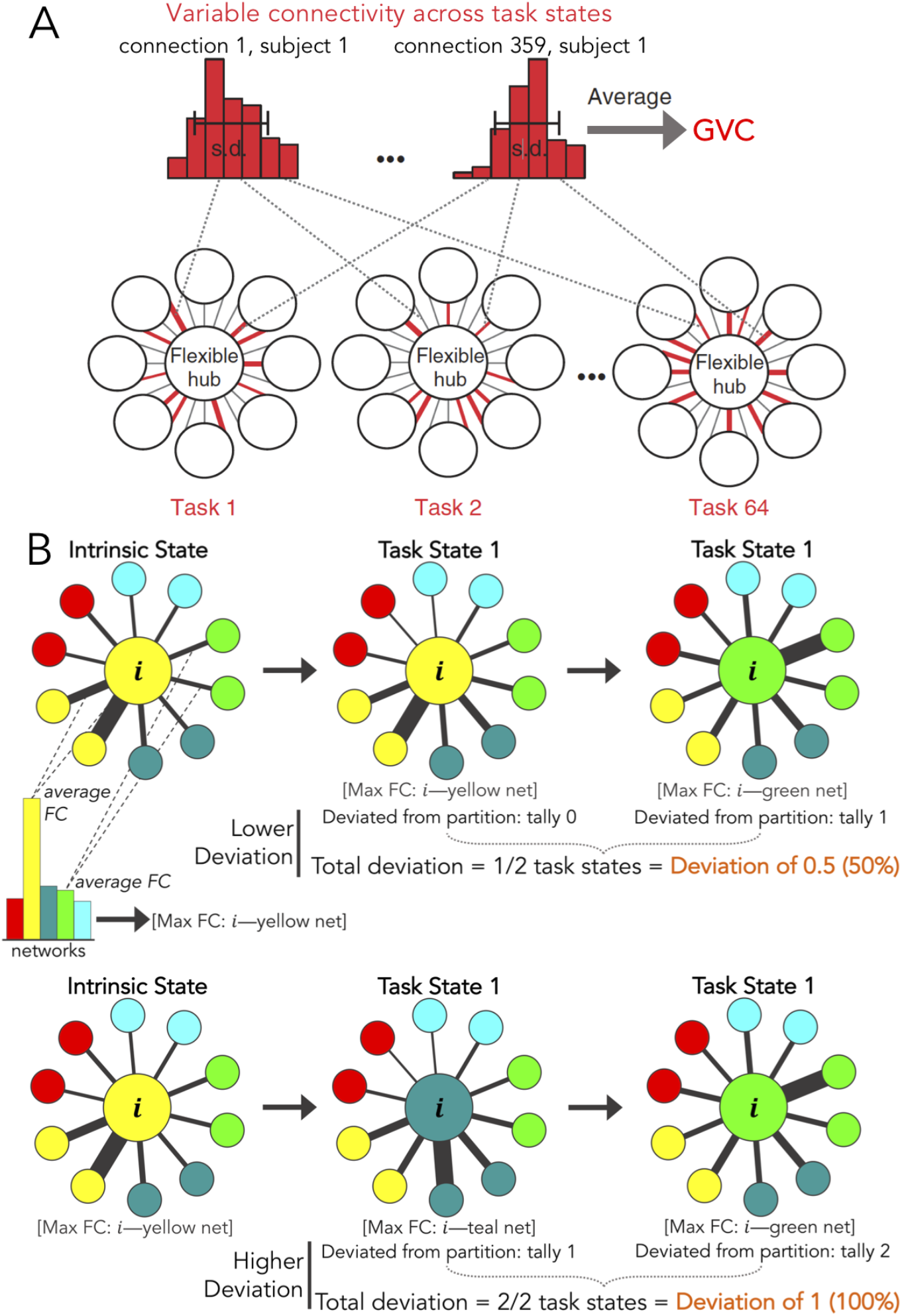
Schematic depictions of network metric algorithms. (***A***) Global variability coefficient (GVC), reproduced with permission (Cole et al., 2013b). Between-network variability coefficient (BVC) is measured equivalently, except within-network connections are withheld. (***B***) Network partition deviation. Per region (large yellow example node labeled “i”): each of its 359 connectivity estimates were averaged according to their CAB-NP (see Materials and Methods and Fig. 3) networks (bar graph in top example), resulting in 12 FC estimates per region. Network “preferences” (network location of maximum FC estimate; thickest lines) were tallied across task states. How often a given region deviated from its predefined partition (intrinsic state) was computed [tally / total number of tasks]. *Lower deviation:* the example region deviated in one out of two hypothetical task states (50% deviation = deviation of 0.5). *Higher deviation:* the example region deviated in two out of two hypothetical states (100% deviation = deviation of 1). The colored nodes encircling the example region represent example regions from example networks, and black lines of variable width represent FC estimates (edge weights).

**Table 3.**
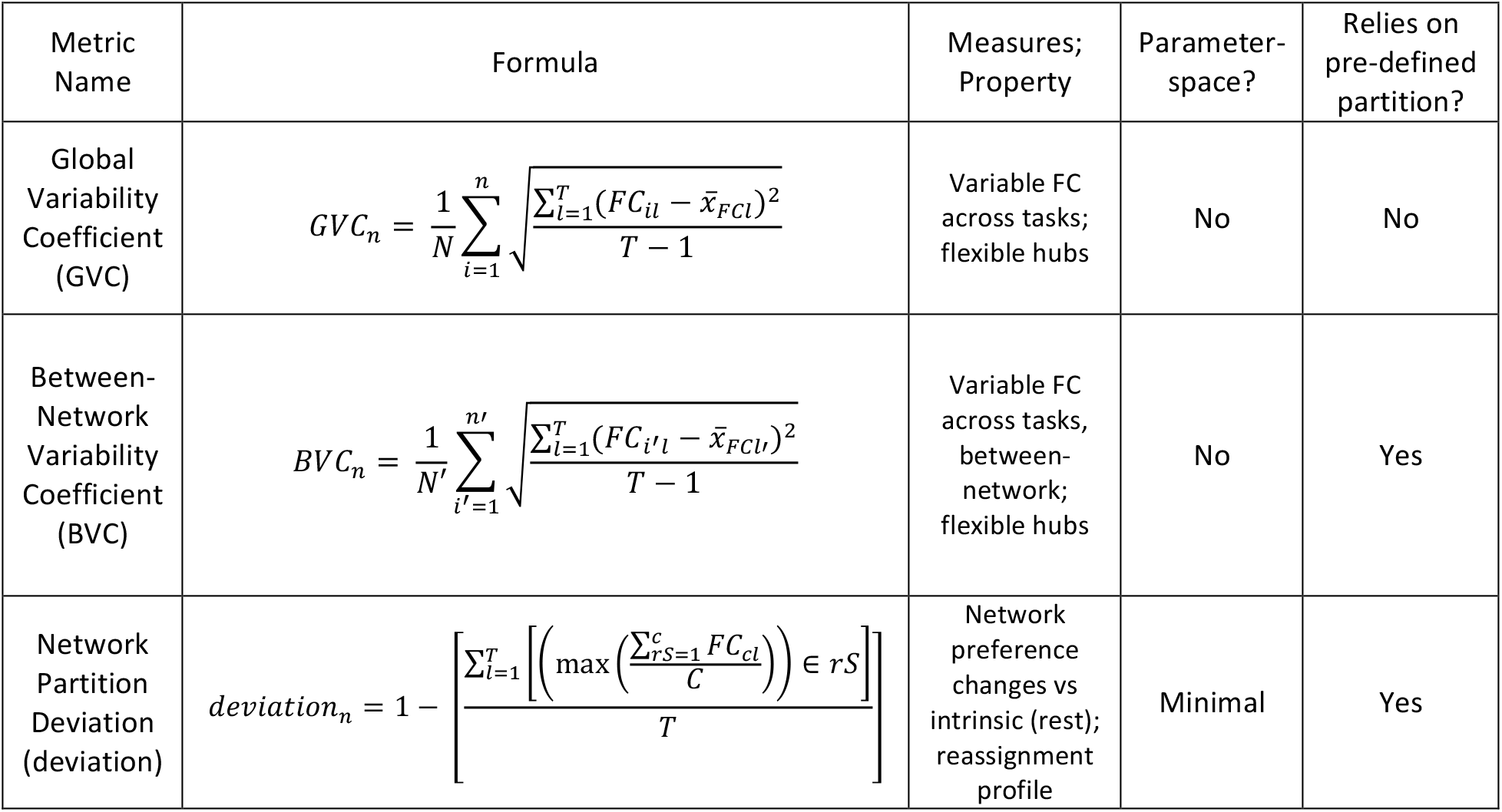
Summary of the network metrics for cognitive control properties across states. Global variability coefficient (GVC; Cole et al., 2013b), between-network variability coefficient (BVC), and network partition deviation (novel) (named in ***column 1***) are described in terms of the following: *(**Column 2**)* their mathematical or algorithmic formulae. All equation symbols are expressed consistently. *Formula terms:* n = brain regions; N = number of regions; i = region 1; l = task 1; T = tasks; FC = weighted adjacency matrix; x-barr = mean; FC_il = edge weight, per region, per task; FCl = FC matrix, per task; n’ = out-of-network regions; N’ = number of out-of-network regions; i’ = region 1, out-of-network; l’ = task 1, out-of-network regions only; FC_i’l = edge weight, per out-of-network region, per task; FCl’ = FC matrix, out-of-network regions only, per task; c = network regions; C = number of network regions; rS = predefined partition; FC_cl = edge weights per network-region, per task. *(**Column 3**)* What each metric measured. This was how results were interpreted, and mechanisms or properties were framed. *(**Column 4**)* If each metric relied on user-chosen parameters (e.g., had a parameter-space). *(**Column 5***) If each metric relied on a predefined network partition (also see Materials and Methods).

### Global Variability Coefficient (GVC)

GVC was originally developed by Cole et al. (2013b), and characterizes changing patterns of connectivity across task states by measuring the variability of interregional connectivity (Fig. 4A and Table 3). Thus, GVC treats spatial changes in connectivity, across states, as continuous. No parameters are required by the user and a predefined network partition is not necessary (aside from regional parcellation, as in the present study) (Table 3).

In Cole et al. (2013b), FPN connections exhibited the highest GVC compared to all other networks. In that study, the FPN also maintained connectivity patterns that could decode task information (using an earlier version of the C-PRO paradigm). Further, FPN connectivity was found to vary systematically with similarity of C-PRO task states. Taken together this suggested that (1) FPN regions exert adaptive task control as flexible hubs, and (2) GVC results were not driven by noise. We replicated these findings and extended the analysis to CON connections. In brief, the 64 C-PRO task states have zero to two overlapping rules (Fig. 1). For example, one task’s rules included *both, high pitch,* and *left middle,* and another included *both, red,* and *left middle.* These example tasks had two overlapping rules *(both* and *left middle).* We created a 64 by 64 similarity matrix to quantify these overlap sets, and quantified the Spearman’s Rho for FPN and CON connections for those sets. Next, we restricted the same analysis by only including FPN and CON regions with the highest GVC, in the following increments: top 10%, 8%, 6%, 4%, and 2%. This addresses whether highly variable connectivity (as measured by GVC) relates systematically to task context (Cole et al., 2013b).

### Between-Network Variability Coefficient (BVC)

BVC was inspired by Cole et al. (2013b), and is related to between-network global connectivity in Ito et al. (2017). BVC is equivalent to GVC, except within-network connectivity estimates are withheld from the computation of standard deviation (Fig. 4A). This change from GVC accounts for the potential confound that within-network connections might confer upon results. In Cole et al. (2013b), FPN had the highest participation coefficient compared to all other networks, suggesting that FPN regions maintain many between-network connector hubs. BVC simply quantifies this in a manner closer to GVC. BVC (unlike GVC) required the use of a predefined network partition to define the regional bounds of each network. All other specifications of BVC are identical to GVC (Table 3).

### Network Flexibility (NF)

NF was originally developed by Bassett et al. (2011, 2013a, 2013b) to quantify how often (i.e., for how many tasks) a region changes its network “allegiance”, and standardizes this by all possible changes. NF is conceptually related to GVC because both metrics quantify large-scale changes in functional network configurations. In the present study, we specifically tested if NF and GVC estimate comparable aspects of network configuration. NF characterizes the spatiotemporal dynamics related to task-state time series by quantifying temporal variability in network partition solutions. These network partitions are determined by an optimized quality function for community detection termed multilayer modularity (also termed multislice or multiplex in some studies) (Louvain Q-modularity; Mucha et al., 2010). Thus, NF does not utilize a predefined network partition, but instead requires community detection to be applied per dataset. Required parameters (g, ω) could be used to tune the degree to which connections were treated as discrete versus continuous in space (g is the spatial resolution parameter) and/or time (ω is the temporal resolution, or coupling, parameter). In the present study, we swept a parameter space of g and ω, ranged around their prototypical values (Bassett et al., 2013; Braun et al., 2015; Chen et al., 2015; Amelio and Tagarelli, 2017). We swept the modularity function’s parameter space by ranging ω from zero to 2.0 in steps of 0.2; and g from zero to 5.0 in steps of 0.5. For both free parameters, zero is the lower limit. An upper limit of 2.0 for ω was based on prior observations that task states tend to merge into one large state at higher coupling values (Bassett et al., 2013a). At the upper limit of g (5.0), spatial resolution becomes acute and each region develops its own network. Additionally, the temporal dynamics conferred by ω are no longer available at the upper limit of g (Chen et al., 2015; Amelio and Tagarelli, 2017). This sweep yielded an 11 by 11 matrix of NF estimates, per region (and per participant). We compared regional NF estimates to regional GVC estimates (both metrics were standardized, and Spearman’s rank-order correlation quantified similarity in these measures across participants), to assess the point in the parameter-space wherein NF and GVC overlap most. These comparisons were performed for both the network-mean and regional-mean vectors. Briefly, we found that NF and GVC characterized shared aspects of network configurations in a specific sector of NF’s parameterspace (see Results). This motivated the development of a novel metric (see next section on network partition deviation) that was less linked to chosen parameters. The remaining Results are based upon GVC and this novel metric (such as the cognitive cartographies; see Results). Importantly, however, there are future research questions that may be better addressed by NF.

### Network Partition Deviation

To reconcile divergent principles and results of GVC and NF (see Results), we created a novel metric termed network partition deviation (or just “deviation”). The primary goal of developing deviation was to quantify network reconfiguration in a highly principled manner. This involved a principled definition for what it means for a region to “reconfigure”: a change in the network community that a given region is most connected with (i.e., the network with the highest mean connectivity). Deviation was the percent of task states (more generally, the relative frequency across time), in which a given region’s “preference” deviated from the predefined partition. To quantify this, deviation enumerated network reassignments from a predefined partition across task states (Table 3 and Fig. 4B). Per task state and per region, connectivity estimates (across the other 359 regions) were searched for the maximum value. The network location of this maximum (relative to the predefined partition) was indexed as the network assignment preference for that given state. To illustrate how network reassignment was computed, we generated a video of the regional network preferences across task states, projected onto a standard brain schematic (Multimedia 1). We used the Connectome Workbench software to generate these visualizations (Marcus et al., 2011).

**Multimedia 1.**
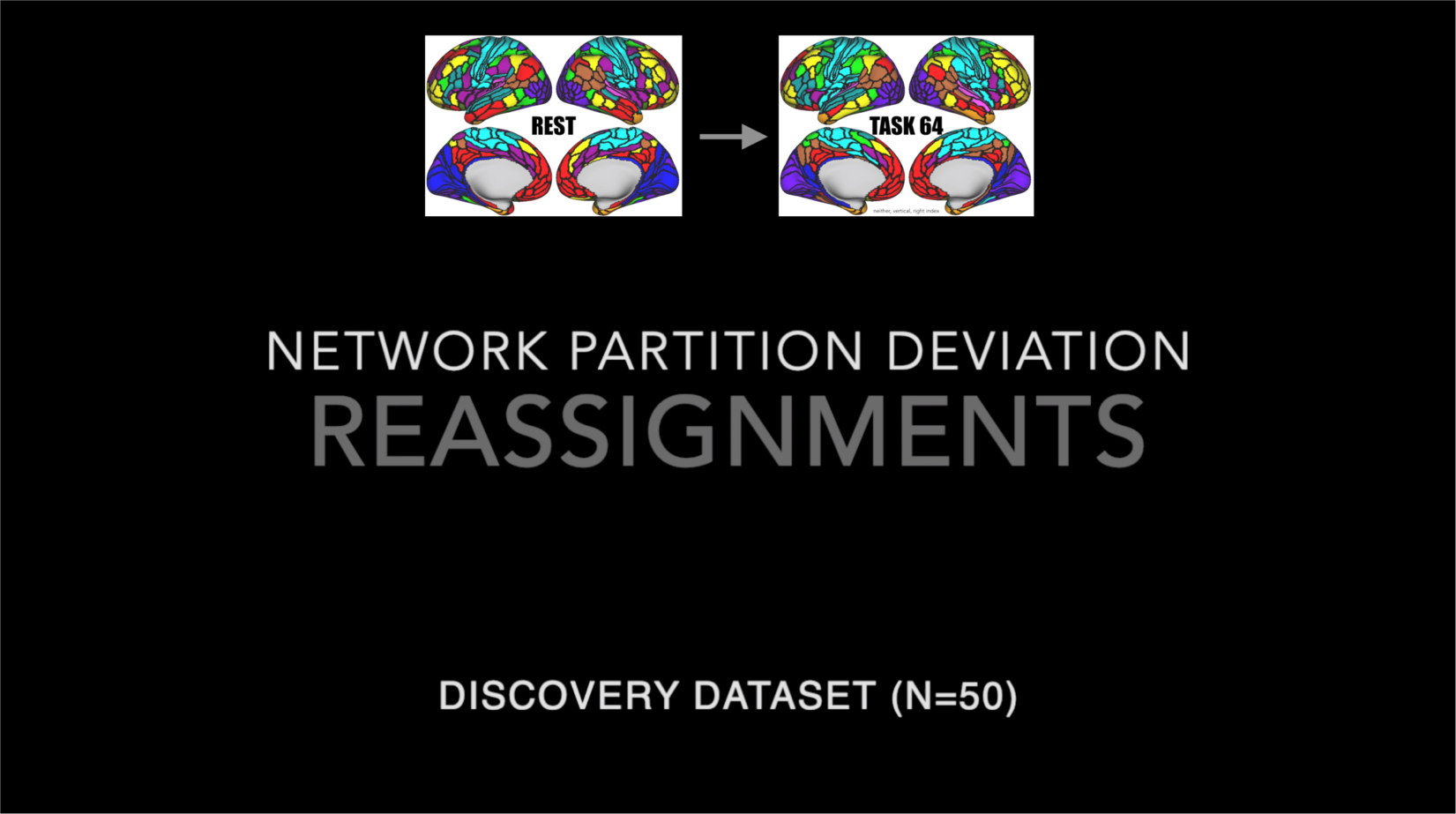
(Preview image) Video depiction of cortical network reassignments across task states, computed via network partition deviation. The video depicts each region’s network “preference” observed during the computation of network partition deviation, across all 64 C-PRO task states, and for *n* = 50 discovery dataset participants (see Materials and Methods). Briefly, per task state and region, the maximum connectivity estimate was found, and its location relative to the predefined partition (empirically adjusted CAB-NP; as in Fig. 3) was indexed (see Fig. 4B and Table 3). These network indices were then mapped back onto brain schematics to visualize how deviation defines network reassignment across tasks. This video also depicts the dynamics captured by deviation. Some patches of cortex remained stable in their network assignments across states (such as primary and secondary visual networks, shown mainly in occipital regions as blue and purple, respectively). However, some regions and/or networks reassigned with more frequency, such as the shifts in cingulo-opercular preference. The goal of deviation was to quantify these patterns in a systematic manner. Cortical regions on each brain schematic represent the Glasser parcellation scheme (2016), and colors correspond to the CAB-NP naming system (Ji et al., 2019; Fig. 3).

We used the Cole-Anticevic brain-wide network partition (Ji et al., 2019), plus adjustments derived from the empirical resting-state fMRI data of the participants studied herein, as the predefined reference (Fig. 3). This *a priori* network partition can be thought of as a minimal parameter space maintained by deviation in the present study, however future work may apply community detection (i.e., empirically-based network partition) if appropriate. Deviation may be accompanied by its complementary measure, network partition adherence, which was the relative frequency of states in which a given region *adhered* to its predefined network assignment (or 1-deviation, meaning that deviation and adherence add up to 1, or 100% of task states). We further unpacked deviation by depicting *which* networks were preferred by regions (when deviating from the partition), generating reassignment profiles.

### Cognitive control cartographies

We rendered one primary mapping of cross-state network-reconfiguration properties, and two secondary mappings which broke up the primary mapping’s properties into within-network and between-network scores. For the secondary mappings, within-network GVC was computed by setting between-network FC estimates to ‘NaN’ (i.e., ‘not a number’ in MATLAB) before inputting data into the GVC algorithm (Fig. 5A). Likewise, between-network GVC was computed by setting within-network FC estimates to ‘NaN’ (Fig. 5B). This effectively nullified the variability for those regions such that GVC ignored them during computation (which principally employs standard deviation across states; Fig. 4A and Table 3). Between-network GVC was equivalent to BVC described above. Within-network deviation was computed by setting FC estimates of between-network regions to the resting-state FC for those regions before inputting data into the deviation algorithm (Fig. 5C). Likewise, for between-network deviation, we substituted within-network estimates with corresponding regions’ resting-state FC (Fig. 5D). We considered the use of resting-state FC most appropriate given deviation’s inherent comparison to the resting-state partition. Thus, deviation away from the resting-state partition would always be zero for regions set to their resting-state estimates. Figure 5 visually depicts the input data schemes for each of these secondary cartographies.

**Figure 5.**
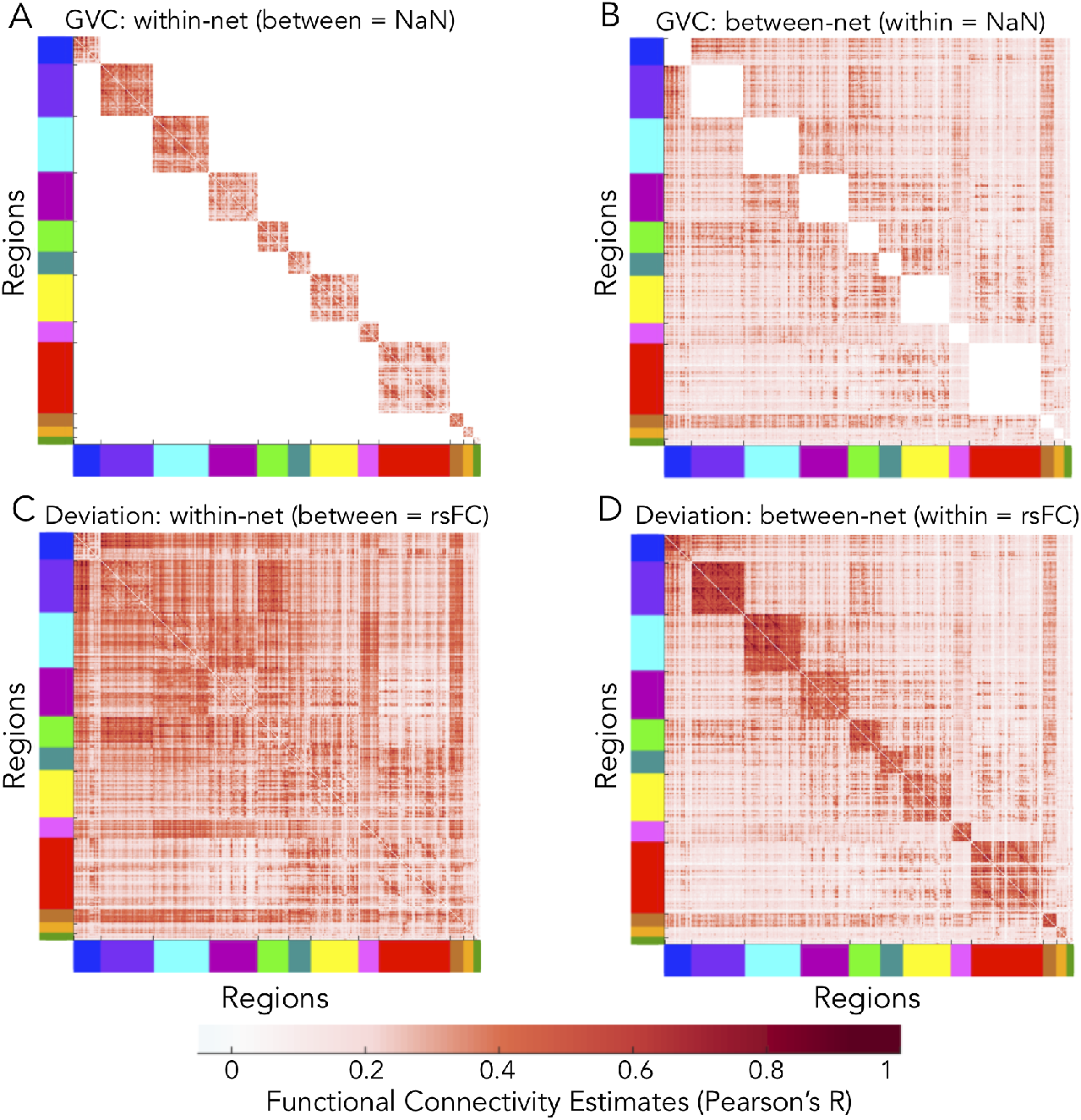
Input data schemes used in computing measures of the secondary cognitive control cartographies. Data refers to functional connectivity estimates (Pearson’s R; colored according to the bottom-most scale). Put another way, each panel contains modified correlation matrices. The cross-state means are visually represented, but analyses included all 64 C-PRO task states. Axes are color coded according to the empirically-adjusted CAB-NP (see Fig. 3). (***A***) Representation of input data for within-network GVC. Between-network FC estimates were set to ‘NaN’ (white). (***B***) Input data for between-network GVC. Within-network FC estimates were set to ‘NaN’ (white). (***C***) Input data for within-network deviation. Between-network FC estimates were set to their resting-state values. (***D***) Input data for between-network deviation. Within-network FC estimates were set to their resting-state values.

### Decoding analyses

A classification analysis was performed for each functional network to test if network connectivity patterns could significantly decode task state. As in Cole et al. (2013b), three 4-way classifications were performed using connectivity patterns from each network separately. We extended the cohort to include *n* = 50 (per discovery and replication datasets), used correlation as a classifying distance measure (Haxby et al., 2001; Mur et al., 2009; Ito et al., 2017), and performed 8-fold cross validation. We performed a within-subjects classification. Each subject had 64 samples of task-state connectivity estimates for each distinct task rule set (see Fig. 1 and Materials and Methods section above on functional connectivity estimation). Of those 64 samples, classifiers were trained on a random subset (over 8 folds) of 56 task states and tested on the remaining (held out) 8 task states. Each task-state was a combination of three rule domains: logic, sensory, and motor. For each of the three decoding analyses, we isolated specific rules from each of these domains. Therefore, the labels associated with these states were according to: (1) logic (both, not both, either, or neither), (2) motor (left middle, left index, right middle, or right index), and (3) sensory (vertical, red color, high pitch, constant tone) rule-set domains (Fig. 1). Therefore, chance accuracy was 25% in each 4-way analysis. We averaged task-state connectivity patterns (i.e., features) across identical training-set labels (e.g., in the logic rule-set classification: training-set connectivity estimates that contained “both” were averaged). We used a minimum-distance classifier (based on Spearman’s correlation score), where a testset would be classified as the rule type whose centroid was closest in the multivariate functional connectivity space (Mur et al., 2009). We compared these distances for each set of matched versus mismatched training and test set labels. When a matched similarity score was larger than all mismatched similarities, this was deemed an accurate decoding. Within-subjects prediction accuracies were equivalent to how many rules were accurately decoded, averaged across 8 folds (Varoquaux et al., 2017).

In order to assess cross-subject statistical significance of the decoding accuracies of each network, we principally performed right-tailed student’s t-tests against chance accuracy. We then utilized the Max-T nonparametric permutation testing approach (1,000 permutations) to address multiple comparisons (see Materials and Methods section above on experimental design and statistical analysis for details). In each permutation, rule-set labels were randomly shuffled before the classification analysis was performed. A null distribution of decoding accuracies and corresponding t-statistics was built and used to assess statistical significance.

### Code and software accessibility

We included all MATLAB, python, and demo code in a publically available platform. Data is available at the level of functional connectivity estimation, for the use of loading into demo scripts. Data at other levels of processing, or data otherwise presented in this study, are available upon request. The master GitHub repository for this study can be found here: https://github.com/ColeLab/controlCartography

## Results

### Intrinsic and Task-State Functional Connectivity

Replicating previous findings (Cole et al., 2014), cortex-wide rsFC (Fig. 6A) and tFC (Fig. 6B) estimates were highly similar. This significant similarity was observed for the average tFC taken across all 64 C-PRO tasks (Mantel-*r* = 0.89, *p* < 0.0001, *R*^2^ = 0.79), as well as for each C-PRO rule individually (Table 4). This aligns with previous observations that the set of networks present during rest are highly related to the set of networks present during task states. In addition to the minimal cognitive demands during rest providing a cognitive baseline for a variety of tasks, this result suggests that rest may be an appropriate intrinsic reference state for characterizing changes in networks across multiple task states. The similarity observed between rsFC and tFC (all 64 C-PRO tasks) in the replication dataset was comparable (Mantel-*r* = 0.90, *p* < 0.0001, *R*^2^ = 0.81).

**Figure 6.**
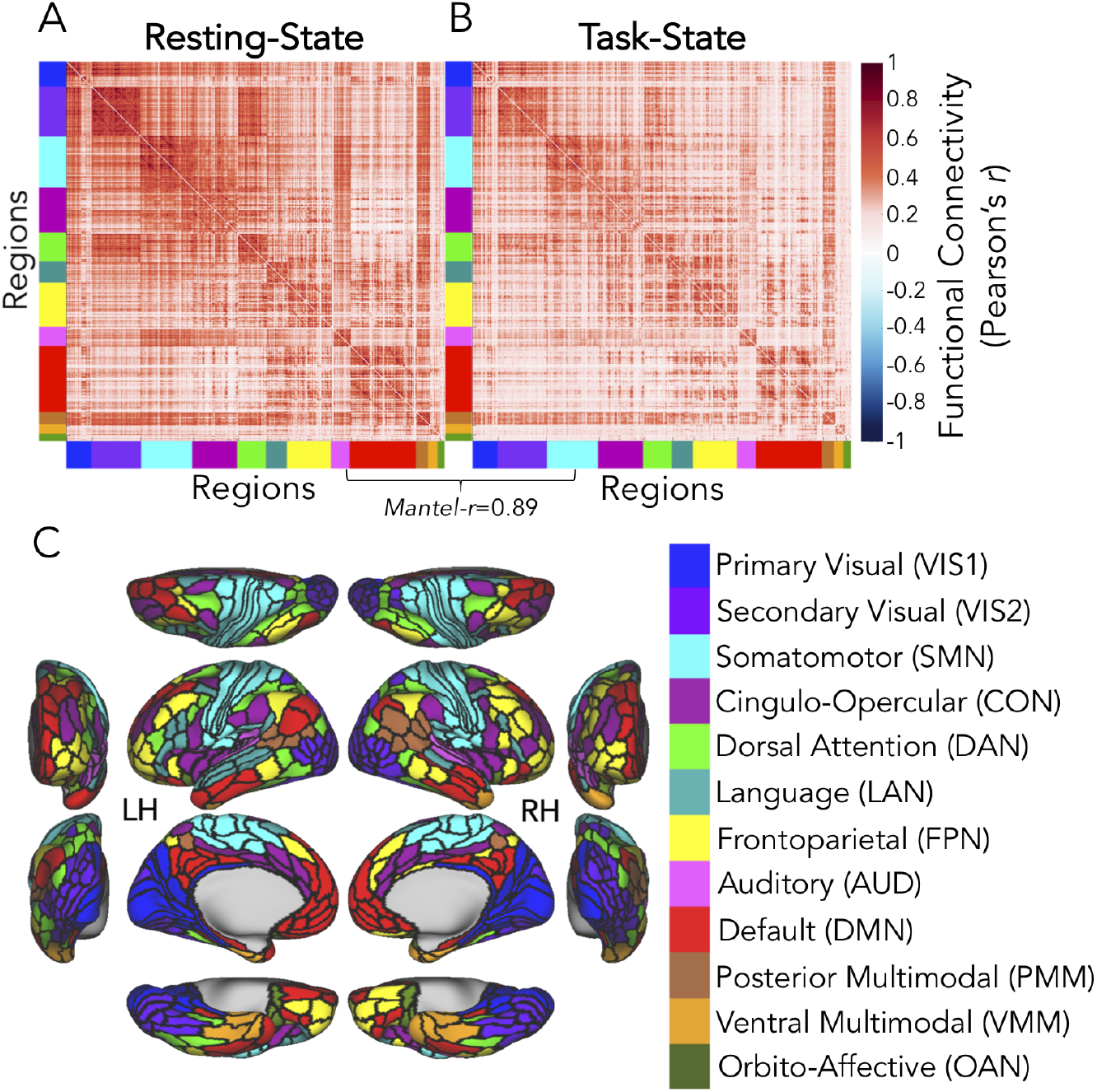
Functional connectivity (FC) estimation. (***A***) Resting-state functional connectivity (rsFC) across 360 by 360 regions (regional parcellation as in Glasser et al., 2016), ordered per the CAB-NP, adjusted by restingstate preferences (see Materials and Methods and Fig. 3) (color-coded along each matrix edge as in C). Discovery-set grand averages are depicted. (***B***) Task-state functional connectivity (tFC) across 360 by 360 brain regions, ordered and estimated as in A (grand averages: *n* = 50 and 64 C-PRO task states). (***C***) Cortical schematic of the Cole-Anticevic Brain-wide Network Partition (CAB-NP) (Ji et al., 2019), empirically adjusted by the resting-state preferences of the present participants (see Materials and Methods and Fig. 3). LH = left hemisphere; RH = right hemisphere. Color-coding scheme of networks and acronyms listed in parentheses are used consistently throughout the present paper. Note that the two cognitive control networks of special interest included the CON (plum) and FPN (yellow).

**Table 4:** Summary of the similarities between intrinsic rsFC and multi-task FC. Each row lists a task-state (i.e., tFC) comparison to rsFC (Fig. 6A). These tFC estimates were based on each of the 12 C-PRO rules (see Fig. 1), and averaged across participants. Columns list the Mantel-r statistic, corresponding *p*-value in scientific notation (nonparametric permutation testing; see Materials and Methods and Glerean et al., 2016), and shared variance (R^2). Connectivity of all C-PRO rule sets significantly correlated with rsFC.

| Table 4. Summary of the similarities between intrinsic rsFC and multi-task FC. |  |  |  |
| --- | --- | --- | --- |
| Task FC Data: C-PRO Rule Set | Correlation with Rest (Mantel-r) | *p*-value (10K permutations) | Shared Variance ($R^2$) |
| Sensory rules |  |  |  |
| Vertical orientation | 0.7966 | $3.05 \times 10^{-8}$ | 0.6345 |
| Red color | 0.7918 | $3.17 \times 10^{-8}$ | 0.6269 |
| High pitch sound | 0.7964 | $3.10 \times 10^{-8}$ | 0.6343 |
| Constant tone | 0.7864 | $3.24 \times 10^{-8}$ | 0.6185 |
| Motor rules |  |  |  |
| Left index finger | 0.7847 | $3.26 \times 10^{-8}$ | 0.6157 |
| Right index finger | 0.7966 | $3.60 \times 10^{-8}$ | 0.6345 |
| Left middle finger | 0.7939 | $3.14 \times 10^{-8}$ | 0.6302 |
| Right middle finger | 0.7844 | $3.33 \times 10^{-8}$ | 0.6153 |
| Logic rules |  |  |  |
| Both | 0.7983 | $3.04 \times 10^{-8}$ | 0.6372 |
| Either | 0.7934 | $3.74 \times 10^{-8}$ | 0.6295 |
| Not both | 0.7833 | $3.28 \times 10^{-8}$ | 0.6136 |
| Neither | 0.7966 | $4.62 \times 10^{-8}$ | 0.6345 |

To summarize changing connectivity from the resting state to the average task state, we created a task versus rest difference matrix, and found 21% of those values to be significant differences (max-T critical threshold = 5.46, 5,000 permutations). The finding that rsFC and tFC (across multiple task states) are highly correlated, yet the differences between them are nontrivial, justified subsequent analyses of functional reconfigurations between these two kinds of states. Findings were consistent in the replication dataset: approximately 32% significant rest-to-task differences (max-T critical = 5.67).

### Network Metrics: Variability Coefficients

Our prior work found that the FPN contains *flexible hub regions* – network nodes capable of rapid reconfiguration with changing task demands (flexible) that have extensive connectivity (hubs) (Cole et al., 2013b). Recent work has suggested that the CON also contains hub-like regions (Ito et al., 2017; Power et al., 2013), yet it is unknown if they are likewise flexible. Accordingly, we used two related metrics to assess if networks contain flexible hub regions (Cole et al., 2013b): global variability coefficient (GVC), and between-network variability coefficient (BVC) (Fig. 4A and Table 3).

Critically, Cole et al. (2013b) only involved *N* = 15 subjects, compared to the *n* = 50 discovery and separate *n* = 50 replication datasets in the present study. Thus, replicating the results of Cole et al. (2013b) would be nontrivial. Replicating the main result of Cole et al. (2013b), regions of the FPN had the highest GVC (Fig. 7A) and BVC compared to the mean of all other networks (GVC: max-T(49) = 10.94, *p* < 0.0001; BVC: max-T(49) = 10.69, *p* < 0.0001). BVC and GVC results were highly correlated at both the network (Fig. 7A) and regional levels (Fig. 7B) (network-wise: *r* = 0.9912, *p* < 0.00001, *R*^2^ = 0.9824 cross-network shared variance; region-wise: *r* = 0.9972, *p* < 0.0001, *R*^2^ = 0.9944 cross-region shared variance), suggesting that within-network estimates do not dominate the outcome of GVC analyses.

**Figure 7.**
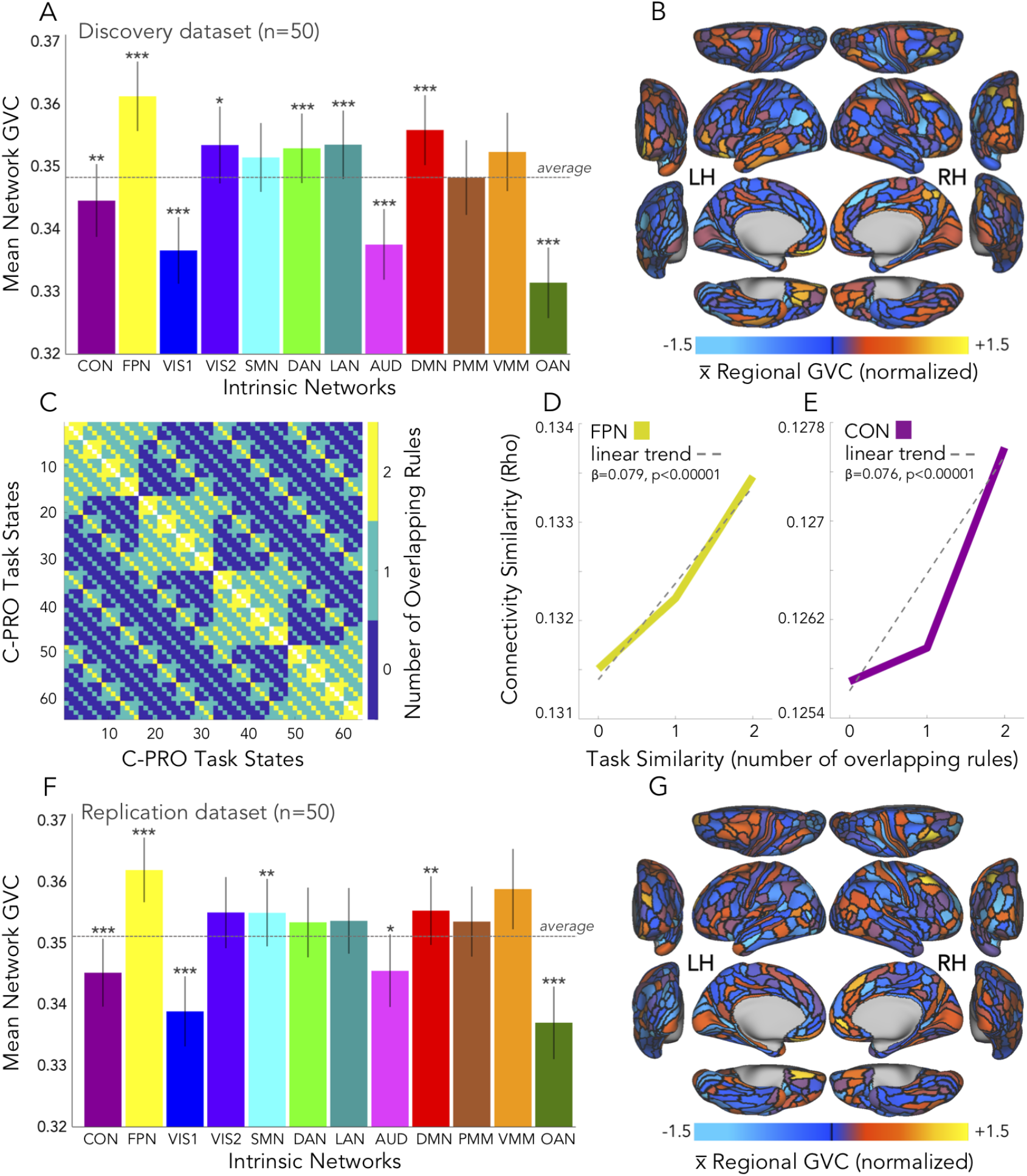
Global variability coefficient (GVC). (***A***) Network-mean GVC, across discovery dataset participants and all C-PRO task states. Error bars: standard error of the mean. Asterisks: statistically significant t-tests, using the max-T approach (see Materials and Methods). Horizontal, dashed line: average GVC across networks. (***B***) Regional-mean GVC, projected onto a cortical surface. LH = left hemisphere, RH = right hemisphere. (***C***) Similarity of C-PRO task states, represented by number of overlapping rules (0 overlapping rule = blue, 1 overlapping rule = green, 2 overlapping rules = yellow). An overlap of 3 rules exists along the diagonal (white), but these connections were not included in analyses because connectivity similarity would be *Rho* = 1 (identical task states). (***D***) Relationship between FPN connectivity similarity and task similarity. All connectivity estimates were included. Grey dashed line: linear trend, with associated beta and t-test significance listed. *(**E**)* Same as D, but for CON regions. The results in D and E demonstrated that control network connectivity similarity varied systematically with task similarity, suggesting that GVC results (A and B) were not driven by network noise. *(**F**)* Same as A, but replication dataset GVC results at the network level. *(**G**)* Same as B, but replication dataset GVC results at the regional level. GVC results highly overlapped between discovery (A) and replication (F) datasets: *Rho* = 0.9091, *p* < 0.00001.

These results were replicated in the replication dataset: the FPN demonstrated the highest GVC (Fig. 7F) and BVC compared to the mean of all other networks (GVC: max-T(49) = 7.23, *p* < 0.0001; BVC: max-T(49) = 6.93, *p* < 0.0001). BVC and GVC results were also tightly correlated at both the network and regional levels in the replication dataset (network-wise: *r* = 0.9925, *p* < 0.00001, *R*^2^ = 0.985 cross-network shared variance; region-wise: *r* = 0.9975, *p* < 0.0001, *R*^2^ = 0.995 cross-region shared variance). Additionally, both GVC and BVC results highly overlapped between discovery and replication datasets (GVC: *Rho* = 0.9091, *p* < 0.00001; BVC: *Rho* = 0.881, *p* = 0.0002).

In conjunction with many studies reporting increased FPN activity as a function of cognitive control demands (Yeo et al., 2015), this pattern of results supports the notion that the FPN contains flexible regions adaptively configured for multitask control. Further, we compared GVC between control networks and each of the other networks. FPN regions were significantly higher than each other network (using the max-T approach, *p* < 0.0001), except for the ventral multimodal network. CON regions were significantly different (typically lower) than each other network on the measure of GVC (using the max-T approach, *p* < 0.0001), except for posterior multimodal and ventral multimodal networks. Lastly, a paired-samples t-test comparing FPN and CON revealed a significant difference in GVC scores (*t*(49) = 11.68, *p* < 0.00001), suggesting that the two proposed control networks exhibit distinct variability of global connectivity. In the replication dataset, FPN regions’ GVC scores were also significantly higher than each other network, except for the ventral multimodal network (orange bar in Fig. 7F). CON regions were significantly different from each other network on the measure of GVC in the replication dataset, with no exceptions (Fig. 7F). The paired-samples t-test contrasting FPN and CON specifically also showed a significant difference on GVC scores in the replication dataset (*t*(49) = 10.55, *p* < 0.00001).

Despite evidence that FPN has strong global variability consistent with flexible hubs, it remains unclear if that variability is systematically related to task information content – a prerequisite for flexible hubs to implement task-related reconfigurations. Prior findings (Cole et al., 2013b) demonstrated that FPN connections systematically vary with increasing task-state similarity. We sought to replicate this result in FPN and – given the current focus on cognitive control systems – we additionally analyzed CON connections. As in Cole et al. (2013b), task-state similarity was taken as the number of overlapping, or shared, rules presented to participants, across all 64 tasks (Fig. 7C; also see Materials and Methods). We then measured Spearman rank correlations (as a score of similarity) amongst connections according to these task-state pairings, for both FPN (Fig. 7D) and CON regions (Fig. 7E). An approximately linear relationship was observed, suggesting that shifts in connectivity systematically relate to shifts in task state, and are not simply a byproduct of noise. Note that at the subject level, the effect size of shifting connectivity is not interpretable because it is unknown how many connection changes are required to cognitively implement a task-rule change (e.g., two robust connection changes may be enough cognitively, but produce small correlation changes at the network level). The linear regression weights of these similarity scores were consistently different from zero across subjects (FPN: *t*(49) = 35.51, *p* < 0.00001; CON: *t*(49) = 33.25, *p* < 0.00001). Next, we performed the same analyses, but restricted the connections to those maintaining the highest variability (across top 2% to 10% in steps of 2%) across task states (i.e., the “VC” of GVC) for both the FPN (as in Cole et al., 2013b) and CON. Results were similar to the main results across all thresholds, with linear weights significantly different from zero (*p* < 0.05). These results suggest that GVC results are likewise driven by systematic changes in connectivity, and not network noise. These results additionally reveal that CON also systematically changes its global connectivity pattern according to task goals, though the GVC results suggest these systematic changes are smaller in CON (and most other networks) than FPN.

Next, we tested the hypothesis that global FPN and CON connectivity patterns were specific enough to each task set that they could be used to reliably predict the current task rules being used. As in Cole et al. (2013b), FPN (as well as all other networks in the present study) features were restricted by their somatomotor network (SMN) connections in the tests of motor-domain rule classification. We tested how well control network connectivity patterns could be used to decode rule sets in the three C-PRO domains (logic, sensory, and motor; Fig. 1) by assessing task decodability of every CAB-NP network (Fig. 3) with nonparametric permutation testing to address multiple comparisons (see Materials and Methods). In each domain there were four distinct rules, thus chance accuracy was 25%.

Consistent with our hypothesis that FPN and CON are especially important for networklevel representation of task information, FPN and CON were the only two networks whose connectivity patterns could be used to decode all three rule domains across both discovery and replication datasets (Fig. 8) (*p* < 0.05, nonparametrically corrected for multiple comparisons). The connectivity patterns of other networks could be used to decode task information in a more limited manner, often for functionally-relevant task domains (e.g., SMN for motor rules; Fig. 8A and Fig. 8C). To clarify the pattern of task-rule decoding results across control and non-control networks, we generated binarized matrices depicting statistical significance (Fig. 8B and 8D). This allowed us to more easily observe which networks’ cortical connectivity patterns could be used to significantly decode rule sets across all three C-PRO task domains (marked by arrows in Fig. 8B and 8D). In the discovery dataset, the cognitive control networks, FPN and CON, plus LAN were the only networks that decode all rule sets. In the replication dataset, only the FPN and CON could significantly decode all rule sets. It is worth noting that LAN came close to maintaining decodability across all rule types in the replication dataset as well, but did not survive correction for multiple comparisons in the sensory domain (*t*(49) = 1.48, *p* = 0.08). Decodability of task information in the language network is consistent with all of the C-PRO rules having been cued with words (Fig. 1). The tendency for control networks’ global connectivity patterns to so consistently carry task rule information in all three domains suggests that their network interactions likely carry information critical to task representation.

**Figure 8.**
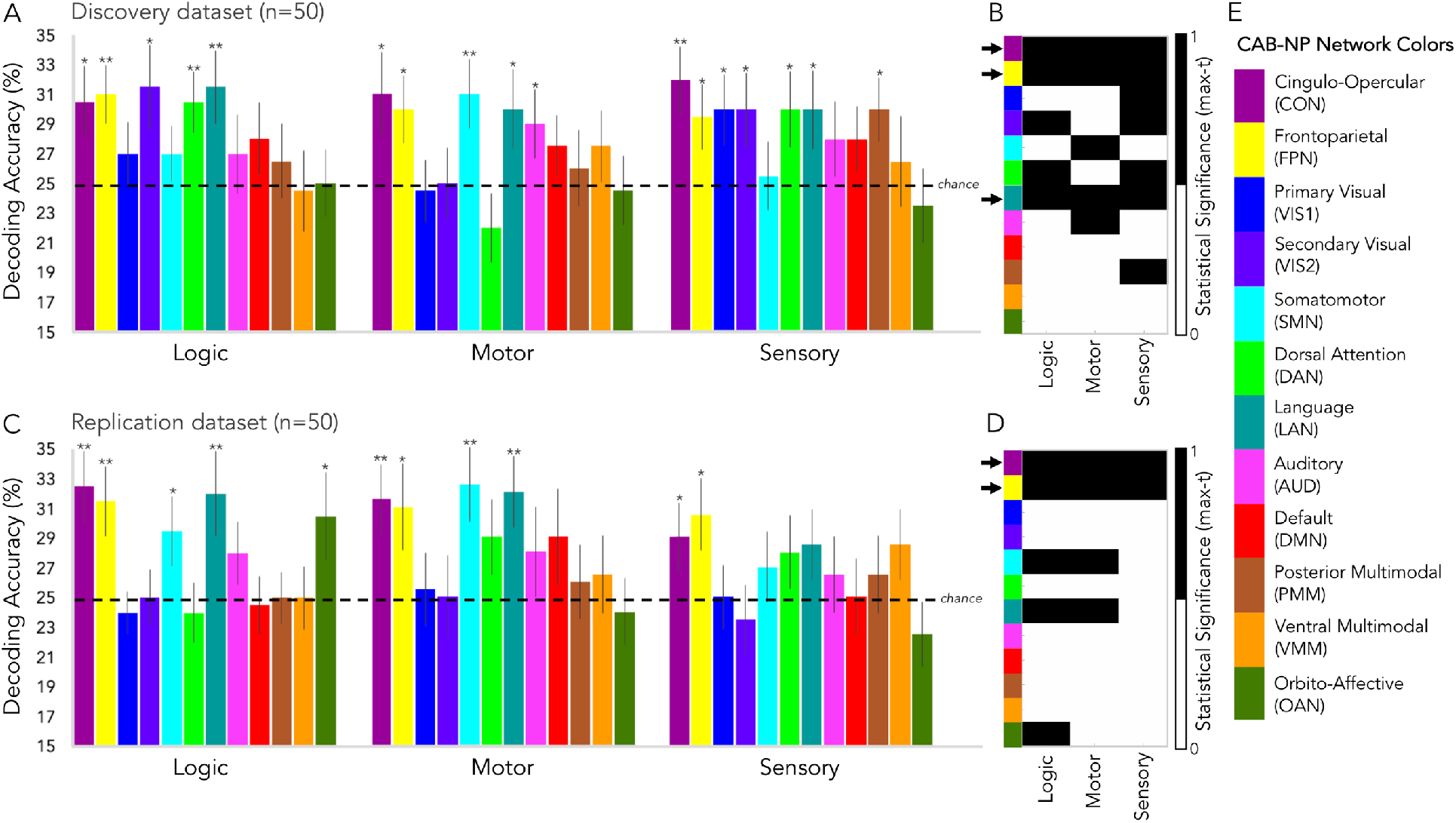
Decoding task rule information with task-state connectivity. (***A***) Cross-subject (discovery dataset) network-mean decoding accuracies for logic, motor, and sensory rules (4-way classifications where chance accuracy was 25%, represented by the horizontal, dashed line). Error bars: standard error of the mean. Asterisks: statistically significant t-tests using the max-T approach (see Materials and Methods). (***B***) Statistical significance tallies (binarized: black = significant or 1, white = not significant or 0) for each network (y-axis, color-labeled) and each rule set (x-axis) for the discovery dataset. Black arrows to the left of y-axis color labels mark networks that significantly decoded all three types of rules, which included the control networks, CON and FPN, as well as the language network (LAN). (***C***) Same as in A, but for the replication dataset. (***D***) Same as in B, but for the replication dataset. The control networks, CON and FPN, were the only networks to significantly decode all three rule types in the replication dataset. LAN came close (as in B), but it’s statistical significance did not survive the permutation testing procedure for sensory rules. (***E***) The CAB-NP color scheme used across all panels A-D (as in Fig. 3, but rearranged to highlight the cognitive control networks). Both FPN and CON connectivity patterns significantly decoded task rules above chance, demonstrating their importance in task representation.

### Network Metric: Flexibility

Network Flexibility (NF) measures functional network dynamics related to task-state time series (Bassett et al., 2011, 2013a, 2013b; Mucha et al., 2010; Cole et al., 2014), and is thus highly relevant to our current hypotheses regarding control network reconfigurations. Conceptually, NF is aligned with GVC, particularly as both quantify large-scale changes in functional network configurations. However, it is unknown whether these metrics capture the same aspects of network reconfiguration. The computations of both GVC and NF are oriented around a measure of network change, however, the approaches are distinct enough to hypothesize that NF and GVC will not entirely overlap. We hypothesized that differentiation between NF and GVC would lend insight into the nature of control network reconfiguration. In particular, GVC assesses continuous changes in connectivity strengths, while NF assesses discrete network reassignments.

The multilayer modularity step required parameters ω (temporal resolution or “coupling”) and g (spatial resolution) to be chosen. The standard values used for these parameters across multiple studies are g = 1 and ω = 1 (Bassett et al., 2013b; Chen et al., 2015; Braun et al., 2015). NF that resulted from community detection at g = 1 and ω = 1 was termed NF-standard. Since there is only a limited theoretical basis for those parameter choices, we computed NF across a range of values around these standards, such that g was varied between zero and five in steps of 0.5; and ω was varied between zero and two in steps of 0.2 (Fig. 9A and 9B). These sweeps resulted in 121 vectors of regional NF estimates (per participant). It was clear that results depended substantially on the exact values of g and ω, such that we were unable to make systematic inferences regarding flexibility of network assignments using NF. To illustrate this: we identified parameters (g = 2.5 and ω = 0.2) that yielded high cross-node similarity to GVC, termed NF-matched (discovery dataset: *Rho* = 0.8169, *p* < 0.00001 as in Fig. 9D; replication dataset: *Rho* = 0.6993, *p* = 0.015) and others g = 0.5 and ω = 0.2 that yielded a negative relationship with GVC, termed NF-unmatched (discovery dataset: *Rho* = −0.4546, *p* = 0.14 as in Fig. 9E; replication dataset: *Rho* = −0.3077, *p* = 0.34), while the NF-standard parameters (g = 1 and ω = 1) yielded a positive but not significant relationship with GVC (discovery dataset: *Rho* = 0.4825, *p* = 0.12 as in Fig. 9C; replication dataset: *Rho* = 0.3147, *p* = 0.32).

**Figure 9.**
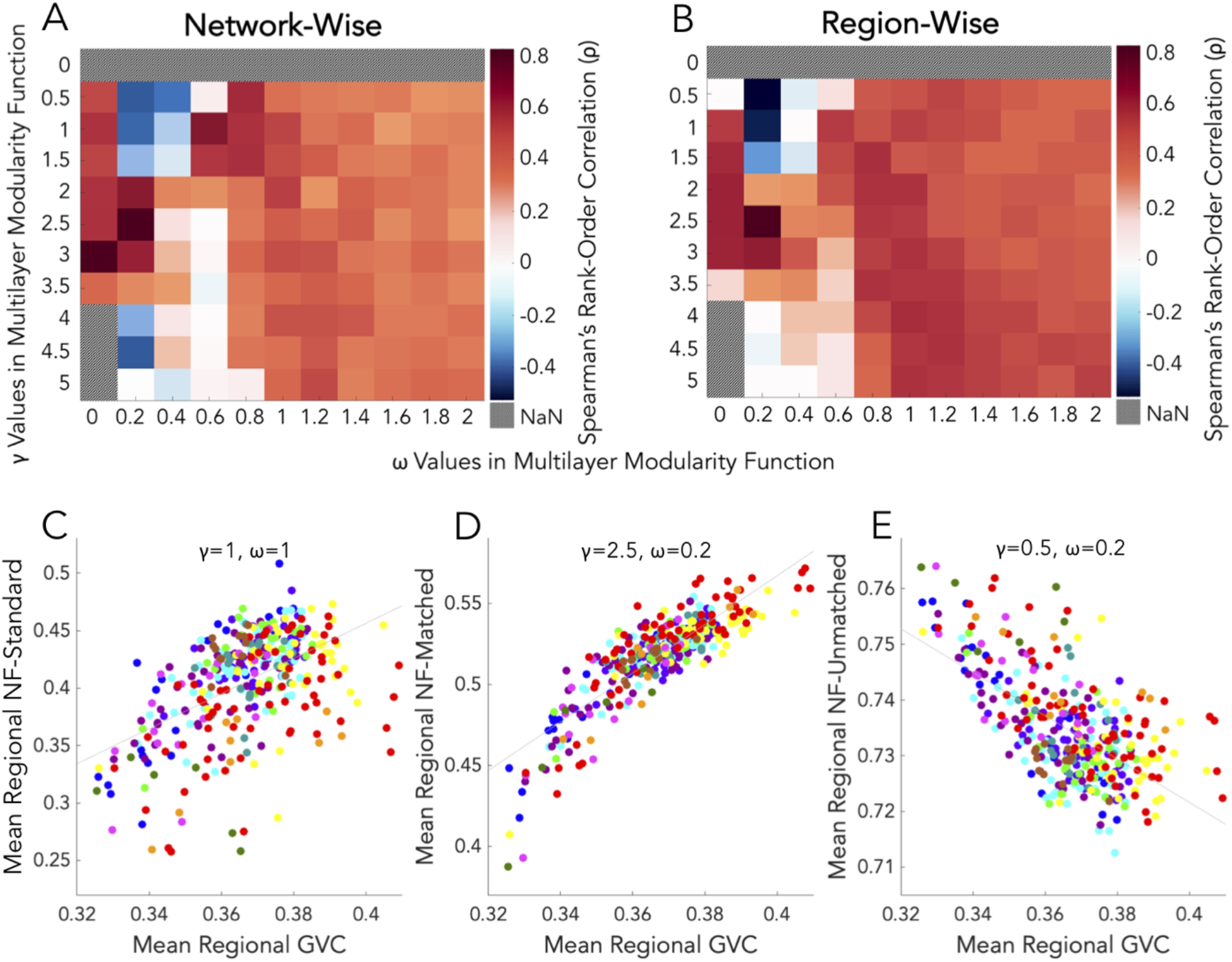
Comparison of global variability coefficient (GVC) and network flexibility (NF), discovery dataset. (***A***) Network-wise comparisons (Spearman’s rank order correlation) of GVC and NF sweeped by multilayer modularity parameters, across participants. (***B***) Same as A, but at the region-wise level. (***C***) Regional mean GVC plotted over regional mean NF-standard (multilayer modularity parameters: g = 1 and ω = 1). NF-standard results were yielded by the standard parameter combination. Each region (mean across *n* = 50 participants) was plotted as individual scatter points, and color-coded according to the network it belongs to (as in Fig. 3). *(**D**)* Same as C, but with NF-matched represented on the y-axis (g = 2.5 and ω = 0.2). This parameter combination yielded NF results most correlated with GVC (or, “matched”), as depicted in A and B by dark red squares. (***E***) Same as C and D, but with NF-unmatched represented on the y-axis (g = 0.5 and ω = 0.2). This parameter combination yielded NF results least correlated with GVC (or, “unmatched”), as depicted in A and B by blue squares. The variable results in panels C-E motivated the need for a new measure of network reconfiguration that was less linked to parameter selection.

### Network Metric: Partition Deviation

Given that NF demonstrated inconsistent results dependent on tuning parameters, we created a novel metric, Network Partition Deviation (or simply, deviation), that could quantify network reconfiguration without the need for a parameter search. Deviation enumerates network reassignments across task states by quantifying the percent of states (i.e., the relative frequency across tasks) in which a given region deviates from a predefined partition (see Materials and Methods, Fig. 4B, Table 3, and Multimedia 1). We employed the CAB-NP adjusted by the empirical resting-state data of participants herein (Fig. 3) as the intrinsic, predefined reference. See prior work (Cole et al., 2014; Krienen et al., 2014) and the Results section “Intrinsic and Task-State Connectivity” above for evidence that resting state provides an appropriate intrinsic network configuration to act as a reference for assessing network deviations.

Of the cognitive control networks of interest here – the CON and the FPN – the CON displayed the highest mean deviation, which was significantly higher than the mean across all other networks (max-T(49) = 12.74, *p* < 0.0001) (Fig. 10A). Moreover, the FPN demonstrated deviation that was near the mean, and was not significantly different from the mean across all other networks. This contrasted from the conclusions drawn from GVC, which showed the FPN significantly above the mean, and the CON significantly below it. Similarly in the replication dataset, CON regions’ deviation scores were again significantly higher than the mean of all other networks (max-T(49) = 16.33, *p* < 0.0001) (Fig. 10E).

**Figure 10:**
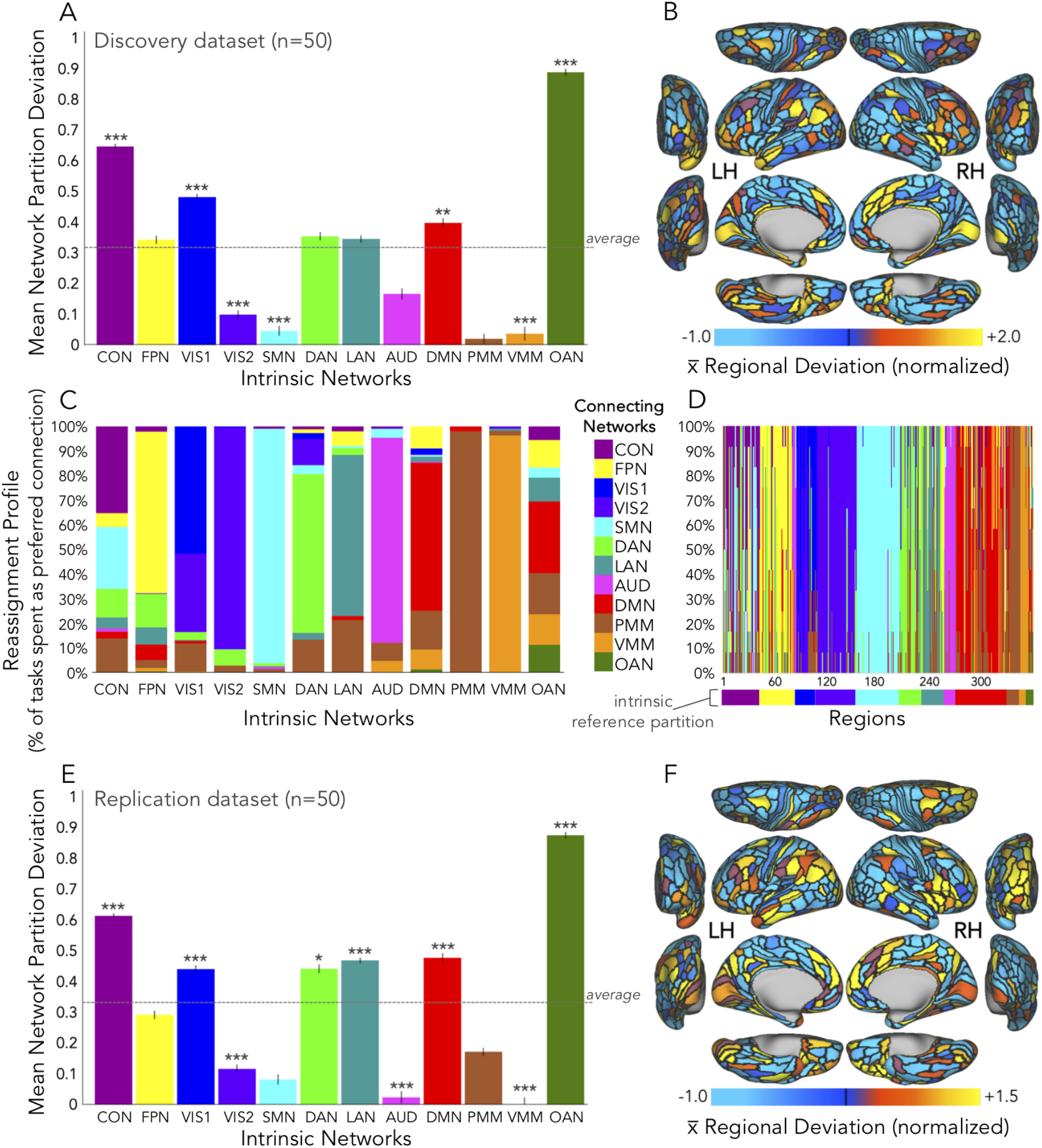
Network Partition Deviation. Mean, error bars, and hypothesis testing specifications were the same as in Fig. 7. (***A***) Network-mean deviation, discovery dataset. See the Discussion for interpretations of the orbito-affective network (OAN). (***B***) Regional-mean deviation, discovery dataset. (***C***) Network-level reassignment profiles. For each intrinsic network (x-axis), adherence (or 1-deviation) is depicted as the portion of the bar (connecting network) with the equivalent color. All other colors codify exactly which connecting networks were being preferred (see Table 3 and Fig. 4) when deviating from the predefined partition. That is to say, deviation in A is expanded in C to show frequency of reassignments, across task states. (***D***) The same as in C, but the x-axis is depicted at the regional level (i.e., these regional reassignments were averaged to generate C). (***E***) Network-mean deviation, replication dataset. (***F***) Regional-mean deviation, replication dataset. Deviation results highly overlapped between discovery (A) and replication (E) datasets: *Rho* = 0.9580, *p* < 0.00001.

Next, we performed planned contrasts of the control networks’ deviation scores versus each other networks’ deviation scores, using the max-T method to correct for multiple comparisons (see Materials and Methods). The CON’s deviation was significantly higher than every other network (*p* < 0.0001), except for the orbito-affective network. Deviation estimates of FPN regions were significantly different from about half of the other networks (*p* < 0.0001), including: secondary visual, somatomotor, cingulo-opercular, auditory, posterior multimodal, ventral multimodal, and orbito-affective. The deviation of FPN and CON regions were significantly different (*t*(49) = 6.28, *p* < 0.00001), suggesting that the control networks differ on how often they deviate from their intrinsic partitions across task states. In the replication dataset, CON regions’ deviations scores were significantly higher than each other network, except for the orbito-affective network. Lastly, the paired samples t-test to compare the deviation of CON and FPN in the replication dataset likewise showed a significant difference (*t*(49) = 9.13, *p* < 0.00001).

To further explore the task-state reconfiguration property that deviation was capturing, we generated “reassignment profiles” at both the network (Fig. 10C) and region (Fig. 10D) levels. Reassignment profiles showed precisely *which* networks were preferred when a region was deviating from the intrinsic partition. As shown in Fig. 10C, the CON deviated to many other networks in an evenly-distributed manner (relative to other networks’ reconfigurations) with some bias to somatomotor connections, whereas the FPN deviated less overall and with more specific preferences, favoring the dorsal attention, language, and default networks (in addition to itself). As with other graph metric results, deviation estimates highly overlapped between discovery and replication datasets (*Rho* = 0.958, *p* < 0.00001), as did reassignment profiles (*Rho* = 0.848, *p* < 0.0001).

### Network Cartographies

We found that FPN regions expressed high GVC yet relatively low deviation. In contrast, CON regions displayed lower GVC yet higher deviation. Altogether, the CON and the FPN both exhibited higher reconfiguration properties than other networks. However, the diversity of findings across network metrics suggested that composite, multidimensional profiles were warranted to fully map out their functionalities. Careful examination of network metric estimates for CON and FPN regions clarified the pattern of results. FPN connectivity tended not to deviate (and when it did, to only a small number of networks as in Fig. 10C and 10D), whereas CON connectivity was more uniform (or “evenly” deviating, as in Fig 10C and 10D). Figure 11A depicts prominent network properties in a cartographic manner (Guimerà and Nunes Amaral, 2005; Mattar et al., 2015), charting GVC on the y-axis against deviation on the x-axis. The FPN can be found in the upper right quadrant of this cartography, near the mean demarcation line for deviation (the vertical gray line, Fig. 11A), pointing to the high-variability yet low-deviation performance of FPN regions in response to cognitive control task state changes. The CON can be found in the lower right quadrant of the cartography in Fig. 11A, suggesting a low-variability yet high-deviation complement to FPN in properties supporting cognitive control.

**Figure 11.**
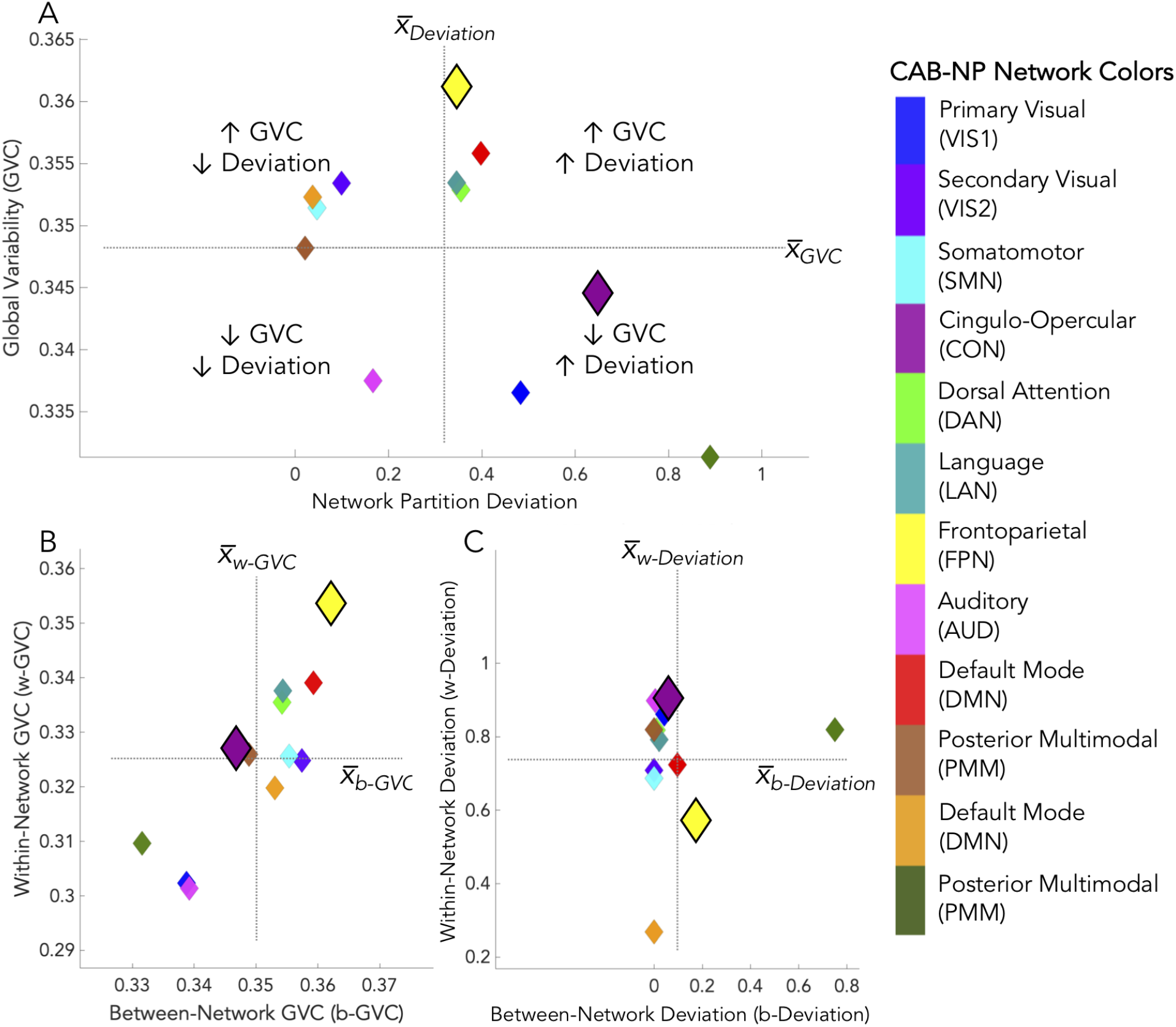
Cognitive control cartographies, discovery dataset. *(**A**)* GVC (as in Fig. 7A) plotted over deviation (as in Fig. 10A), with demarcation lines (dashed and crossed gray lines) indicating the cross-network mean for each dimension (all axes are centered at these marks for ease of viewing). This allowed us to “map out” multidimensional properties at once. For example, networks in the lower right quadrant of A (such as CON) exhibited GVC lower than the mean and deviation higher than the mean. This mapping suggests a nonlinear relationship between GVC and deviation, suggesting that each measure characterized a unique network property. In all panels, control network diamonds (FPN: yellow, CON: plum) are highlighted with dark black outlines and are larger than other network diamonds, with the sole visualization purpose of standing out as cognitive control networks. (***B***) GVC scores (y-axis of panel A) expanded by within-network and between-network values. (***C***) Deviation (x-axis of panel A) scores expanded by within-network and between-network values. The far-right legend depicts the CAB-NP color scheme (as in Fig. 3) used for the diamonds.

To expand upon the primary mapping in Fig. 11A, we generated two secondary cartographies that depict the quantification of each primary measure’s within-network and between-network scores (see Materials and Methods). Briefly, within-network dynamics were assessed by keeping the between-network connectivity fixed across states (defined by resting state). Similarly, between-network dynamics were assessed by keeping the within-network connectivity fixed across states (defined by resting state) (see Fig. 5 for data input schematics). Figure 11B charts these dimensions of GVC, showing both within-network and between-network FPN connections to be high on global variability, and CON to be near-mean on both within-network GVC and between-network GVC. Figure 11C depicts within- and between-network deviation. Both FPN and CON regions were near the mean for between-network deviation (yet on opposing sides of the mean, see vertical gray line in Fig. 11C), yet low and high, respectively, for within-network deviation. This suggests that CON’s high deviation was driven primarily by changes in within-network connectivity.

Supporting dissociation of FPN and CON in terms of network dynamics, we directly compared FPN and CON regions on each secondary cartographic metric (Fig. 11B and 11C), and found the following: (1) FPN was significantly higher than CON on within-network GVC (*t*(49) = 8.92, *p* < 0.00001); (2) FPN was significantly higher than CON on between-network GVC (*t*(49) = 11.43, *p* < 0.00001); (3) CON was significantly higher than FPN on within-network deviation (*t*(49) = 6.55, *p* < 0.00001); and (4) FPN was significantly higher than CON on between-network deviation (*t*(49) = 4.88, *p* = 0.000012) (see prior results sections for FPN versus CON comparisons on GVC and deviation scores related to Fig. 11A, where all regions were included). This pattern of results replicated in the replication dataset: (1) FPN was significantly higher than CON on within-network GVC (*t*(49) = 8.65, *p* < 0.00001); (2) FPN was significantly higher than CON on between-network GVC (*t*(49) = 10.26, *p* < 0.00001); (3) CON was significantly higher than FPN on within-network deviation (*t*(49) = 6.16, *p* < 0.00001); and (4) FPN was significantly higher than CON on between-network deviation (*t*(49) = 2.13, *p* = 0.03).

To explore these results further, we created a color-coded graph of the partition reassignments captured by each version of deviation (Fig. 12). To quantify the relationships between reassignment patterns we used the Jaccard similarity index (see Materials and Methods). We found FPN’s between-network deviation (i.e., within-network connectivity held constant) was more similar to “all-data” deviation than its within-network deviation (i.e., between-network connectivity held constant): Jaccard indices of 0.29 and 0.18, respectively (Fig. 12B). Yet, CON’s within-network deviation was more similar to “all-data” deviation than its between-network deviation (Jaccard indices of 0.31 and 0.16, respectively; Fig. 12A). Supporting dissociation of FPN and CON network dynamics, the Jaccard similarity indices for CON and FPN were significantly different (Jaccard for deviation all-data and deviation within-network data, CON vs FPN: *t*(49) = 3.30, *p* = 0.0018; Jaccard for deviation all-data and deviation between-network data, CON vs FPN: *t*(49) = −4.61, *p* = 0.00003).

**Figure 12.**
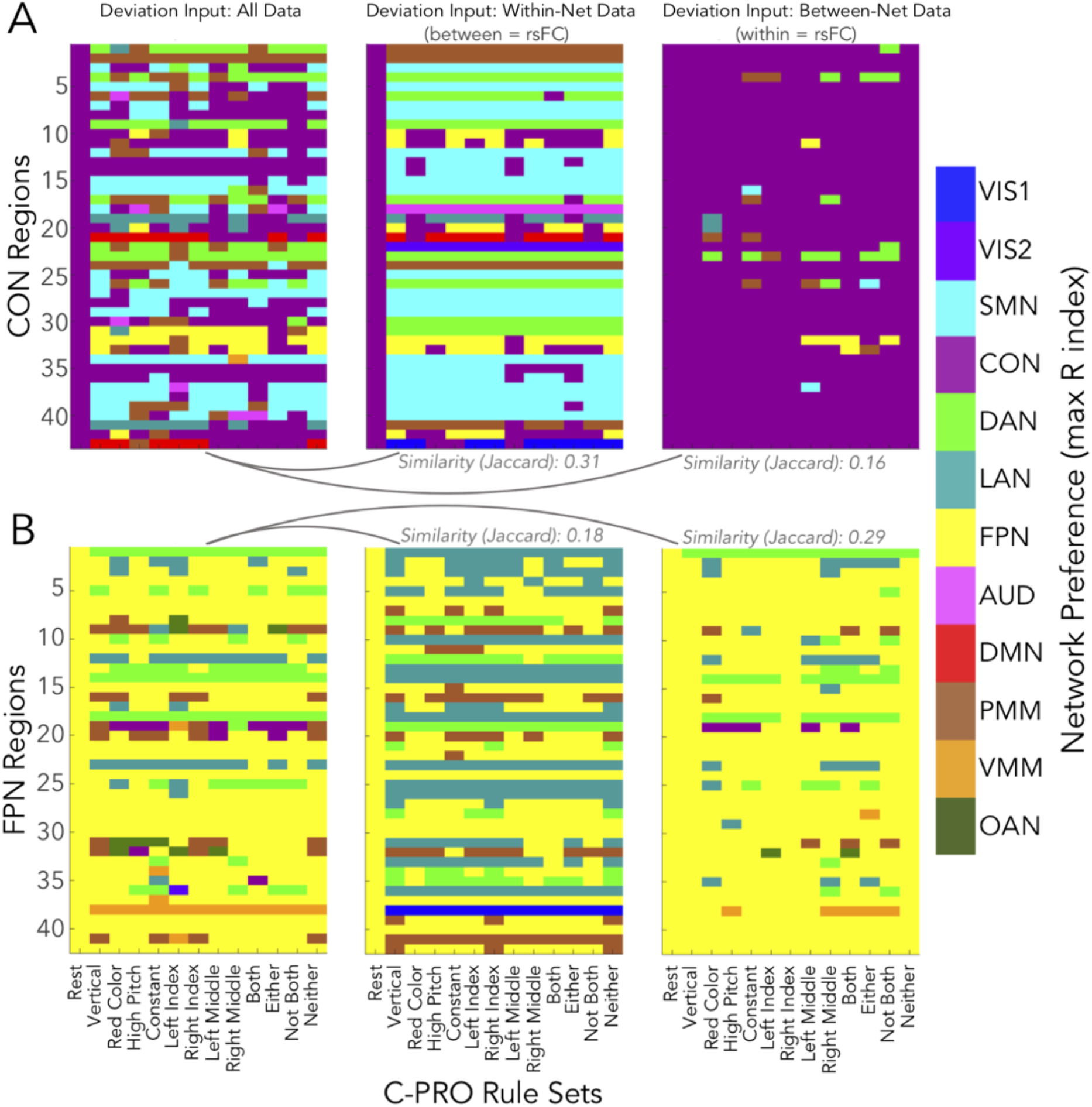
Reassignments conferred by the variants of network partition deviation in the control cartographies. Reassignment is the network index (based on the intrinsic partition) of the highest mean connectivity estimate, per task state. In each panel, Jaccard indices are listed to indicate similarity between two partitions. Network assignments are color coded (as in Fig. 3), and the 64 C-PRO task states are collapsed into 12 rule sets (plus Rest as a reference on each x-axis). (***A***) *Left:* Network reassignments of CON regions from the deviation algorithm, with all connectivity data included in the input (Fig. 11A, x-axis). *Middle:* Within-network CON estimates used in the deviation algorithm (Fig. 11C, y-axis; see Fig. 5C). The Jaccard similarity of within-network and all-data deviation is 0.31. *Right:* Between-network CON estimates used in the deviation algorithm (Fig. 11C, x-axis; see Fig. 5D). The Jaccard similarity of between-network and all-data deviation is 0.16, which is lower than the within-network similarity to all-data. (***B***) Same as panel A, except for FPN regions. The Jaccard similarity scores are: within-to-all = 0.18, between-to-all = 0.29. Thus, FPN showed the reverse pattern to CON, where between-network deviation is more similar to all-data deviation than within-network deviation.

This result supports the conclusion that the high deviation exhibited by the CON was driven by its within-network connections, indicating that CON task-related dynamics were driven mostly by reduction in within-network intrinsic connectivity (network “disbanding”) to increase the strength of between-network connections relative to (now-reduced) within-network connections. In contrast, FPN regions maintained their within-network connection patterns while varying their between-network connection patterns across rest and task. This is consistent with FPN maintaining its intrinsic within-network connectivity while reconfiguring its between-network connections in a task-specific manner. Moreover, the pattern of CON within-network decreases were task-specific (Fig. 7E, Fig. 8, and Fig. 12). This is consistent with CON being a flexible hub network like FPN, but with a distinct mechanism involving “switching” from within-network to out-of-network connectivity via dynamic reduction of within-CON connectivity.

## Discussion

The chief conclusion of the current study was that combining network science measures into network cartographies (multi-dimensional functional “mappings”) allowed us to characterize cognitive control brain systems as either flexible coordinators (frontoparietal regions) or flexible switchers (cingulo-opercular regions). Network cartographies consisted of two primary dimensions: 1) GVC, which measures global FC reconfiguration across task states in a continuous manner and, 2) deviation, which measures global FC reconfiguration from rest to task in a discrete manner (Fig. 11). We found that FPN exhibited high GVC but low deviation, while CON showed the opposite pattern, consistent with complementary mechanisms of cognitive control. FPN appeared to act as a “flexible coordinator”, based on its extensive between-network FC reconfiguration along with maintenance of its within-network connections across rest and task states. In contrast, CON appeared to act as a “flexible switcher”, based on extensively reducing its within-network connections from rest to task to effectively switch to other networks during tasks (Fig. 2).

The present findings are broadly consistent with the view proposed by Dosenbach et al. (2006, 2007, 2008), which posited – based on fMRI task activations and resting-state FC – that control networks implement dissociable mechanisms. We used tFC along with dynamic graph-theoretic measures to expand on Dosenbach et al. In that work, FPN regions enacted control in a manner described as “active, adaptive, and online”. The high tFC-based global variability we observed in FPN regions is consistent with adaptive monitoring and adjustment important for controlled processing (Cole and Schneider, 2007; Sadaghiani and D’Esposito, 2015; Crittenden et al., 2016). In contrast to FPN, Dosenbach et al. (2006, 2007, 2008) proposed that CON underlies “stable set maintenance, task mode, and strategy” (also shown by Vaden et al., 2013). While we did find CON connectivity changes to be more consistent (across task states) than FPN (Fig. 7), we propose that its functional switching relates to the biased competition model put forth by Desimone et al. (1990, 1995) and related theories, as described below.

The biased competition model posits that neural representations compete for resources, such that stimuli, actions, and/or thoughts compete for attention during task performance (Desimone and Duncan, 1995). The theory suggests that competition is biased by top-down goal-related signals from prefrontal cortex and related areas (i.e., control networks). These top-down control signals are thought to shift the competition in bottom-up processing (e.g., in visual cortex), such that goal-relevant processes become more salient and more likely to “win”. For instance, a top-down control signal could bias color-naming representations over word-reading representations to aid in Stroop task performance. This theory was built upon by the guided activation theory (Miller and Cohen, 2001) and flexible hub theory (Cole et al., 2013b). In line with these later theories, we recently posited that such top-down biases to bottom-up competitive processes are especially important for RITL paradigms (such as the C-PRO task), and that they are implemented by tFC changes from control networks (Cole et al., 2017). In the present study, connectivity patterns of FPN and CON regions significantly decoded C-PRO task rules (Fig. 8), suggesting that distributed interactions implemented by cognitive control networks critically support task representations. Further, network science measures probing those interactions suggested that top-down biases are implemented via two complementary mechanisms.

First, CON regions appeared to reduce their within-network connectivity and flexibly switch to other networks in a task-dependent manner. We observed this switching to occur with a relatively uniform distribution, across tasks and switched-to networks (Fig. 7 and Fig. 10C), and with high deviation across tasks (Fig. 10A). We posit that CON transiently disbands and switches networks to lend resources (“weight” or “energy”) to help goal-relevant regions/networks (e.g., visual and motor regions during visuo-motor tasks) win competitions with other regions/networks (or representations). Importantly, we propose that CON’s switching property specifically helps win competitions by reducing functional interference from goal-irrelevant systems, such as interference with distracting stimuli or amongst goal-relevant representations. Second, FPN regions appeared to flexibly coordinate their global patterns of goal-driven biases with each other via maintaining within-network connectivity. This likely facilitates the coordination of complex task sets via facilitating interactions amongst combinations of task representations. This account illustrates a fundamental trade-off in controlled processing: implementing goal-relevant “programs” by FPN through coordinated (but potentially interfering) top-down biases, versus lending of resources via independent (and therefore less likely to interfere) top-down biases by CON to help goal-relevant brain systems win competitions.

Interestingly, FPN’s between-network connectivity patterns were variable, but FPN’s within-network intrinsic configuration remained intact across task states (Fig. 2 and Fig. 12). On the one hand, between-network FC variability corroborates the notion that FPN supports task-specific coding (Crittenden et al., 2016) and selective attention demands (Sadaghiani and D’Esposito, 2015). On the other hand, within-network FC preservation suggests that FPN regions are *coordinating* the FC changes across FPN regions. These dynamics are well-suited to address the “variable binding” problem (Feldman, 2013), where variable stimulus information must be linked to task rules to enact cognitive computations. In C-PRO tasks, variable rules must link via logical operations to perform a given task, and variable stimuli must link to those rules to produce correct behavior (Fig. 1). The maintenance of FPN’s intrinsic organization combined with between-network reconfigurations, suggests a coding process that includes FPN, along with other, task-specific regions. This computational format would allow for variable stimulus information to be bound on a task-to-task basis. Specifically, we propose that FPN’s role in this scheme is to flexibly coordinate task-specific coding. Notably, this would impose high processing demands on FPN regions, which we suggest to be facilitated by the CON freeing up resources, pointing to a computational trade-off across these two cognitive control networks.

Two other networks joined CON in having low variability and high deviation: the OAN and the primary visual network (VIS1). OAN had both the lowest GVC and the highest deviation of all networks. This suggests that the switching mechanism proposed for CON also occurs for OAN, though other studies suggest distinct functionality for OAN. Anatomically, OAN is localized to a small number of regions – ventromedial prefrontal cortex and nearby subcortical regions (Ji et al., 2019). Human lesion studies and animal models suggest a core role of OAN in emotion processing and value representation (Roy et al., 2012), and likely receives direct dopaminergic projections from the ventral tegmental area (Seamans and Yang, 2004). Emotion processing may seem counter to the non-emotional C-PRO paradigm, yet evidence from human lesion studies (Koenigs et al., 2007) and neuroimaging (Botvinick and Braver, 2015) demonstrates that emotion, in the form of motivation, biases competition between outcomes during complex decisions. Future research should assess whether OAN provides a similar mechanism to CON for top-down biasing, but via an emotional/motivational mode of processing. In contrast to OAN and CON, VIS1 was not diverse in its connectivity switches, primarily switching to the secondary visual network (extrastriate cortex) (Fig. 10C and 10D), consistent with integrated processing across the two visual networks.

Aside from the high-variability/low-deviation and low-variability/high-deviation cartographic mappings of the FPN and CON, respectively (Fig. 11A), there were two other scenarios possible. First, high-variability and high-deviation across task states: networks exhibiting this profile would have fluctuating connectivity as well as notable partition reassignment from rest to task. The default network (red diamond in Fig. 11A) appeared to be the only network trending in this direction. Secondly, low-variability and low-deviation across task states: networks exhibiting this profile would have stable connectivity estimates and adhere to their intrinsic partition. The auditory network was the only observed herein (pink diamond in Fig. 11A), which may relate to sensory regions’ proposed “rigid core” organization (Bassett et al., 2013b). Other sensory-motor networks had low deviation compared to cognitive networks, suggesting low vs. high deviation was indicative of sensory-motor vs. cognitive network properties. Future work is warranted to explore this, particularly if diverse task paradigms are implemented.

An essential consideration for future studies regards the question of timescale. Dosenbach et al. (2007, 2008) found increases in CON activity to be sustained across tasks, while FPN activations were present at task onset then adaptively varied with changing task demands. In the present work, we applied network metrics across a set of dynamic task states demanding high levels of cognitive control. While measures of global variability summarized varying connectivity patterns across states, further examination could determine the *timescales* of control network mechanisms. Relatedly, future studies would benefit from considering how electrophysiological signatures of neural processing and network properties relate in terms of the instantiation of cognitive control. In resting-state based studies, Sadaghiani et al. (2010, 2012) found distinctions between FPN and CON based on alpha band signatures. Spontaneous CON activity related to increases in global power, while FPN related to increases in long-range phase synchrony. These signatures correspond to the functions of tonic alertness and phasic control, respectively. In future work, both rest and control-related task states should be assessed via alpha band signatures, as well as potential changes in those signatures from rest to task. A potential outcome is that increases in global power (CON) and long-range synchrony (FPN) would be more apparent from rest to task, constituting another “mappable” reconfiguration property of cognitive control. This would further support the proposition that CON is suited to lend processing resources, and FPN to adaptively integrate task-specific information. Moreover, possible interactions between the properties discovered herein (Fig. 11) and electrophysiological properties remains an empirical question.

Taken together, constructing a functional cartography by combining multiple network science measures allowed us to characterize FPN and CON as complementary systems of cognitive control. We demonstrated that FPN regions enacted control via flexible coordination of reconfiguring connectivity patterns, and CON regions enacted control via flexible switching of network affiliations to lend resources to task-relevant networks. All results replicated in a dataset with distinct subjects, and expanded prior theories that distinct mechanisms of cognitive control are instantiated in parallel via separate large-scale brain systems. Looking forward, we expect the dynamic network neuroscience approach expanded upon here will be effective for functionally characterizing the relationship between neural and cognitive dynamics in other brain systems and other cognitive paradigms.

## Acknowledgments

This project was supported by the US National Institutes of Health, under awards K99-R00 MH096901 and R01 MH109520. The content is the sole responsibility of the authors and does not necessarily represent the official views of any of the funding agencies. The authors acknowledge the Office of Advanced Research Computing (OARC) at Rutgers, The State University of New Jersey for providing access to the Amarel cluster and associated research computing resources. The authors thank their colleagues at the Cole Neurocognition Lab and the Rutgers University Brain Imaging Center (RUBIC) for their expertise and diligent efforts in data collection, as well as offering words of wisdom and overall support.

